# Structures of the human cholecystokinin 1 (CCK1) receptor bound to Gs and Gq mimetic proteins: insight into mechanisms of G protein selectivity

**DOI:** 10.1101/2021.05.06.442871

**Authors:** Jesse Mobbs, Matthew J. Belousoff, Kaleeckal G. Harikumar, Sarah J. Piper, Xiaomeng Xu, Sebastian G.B. Furness, Hari Venugopal, Arthur Christopoulos, Radostin Danev, Denise Wootten, David M. Thal, Laurence J. Miller, Patrick M. Sexton

## Abstract

G protein-coupled receptors (GPCRs) are critical regulators of cellular function acting via heterotrimeric G proteins as their primary transducers with individual GPCRs capable of pleiotropic coupling to multiple G proteins. Structural features governing G protein selectivity and promiscuity are currently unclear. Here we used cryo-electron microscopy to determine structures of the CCK1R bound to the CCK peptide agonist, CCK-8 and two distinct transducer proteins, its primary transducer Gq, and the more weakly coupled Gs. As seen with other Gq/11-GPCR complexes, the Gq-α5 helix bound to a relatively narrow pocket in the CCK1R core. Surprisingly, the backbone of the CCK1R and volume of the G protein binding pocket was essentially equivalent when Gs was bound, with the Gs α5 helix displaying a conformation that arises from “unwinding” of the far C-terminal residues, compared to canonically Gs coupled receptors. Thus, integrated changes in the conformations of both the receptor and G protein play critical roles in the promiscuous coupling of individual GPCRs.

**One-Sentence Summary:** Cryo-EM structures of the CCK-1R reveal key mechanisms for promiscuous G protein coupling.

## Main Text

G protein-coupled receptors (GPCRs) are ubiquitous regulators of cellular function, acting as allosteric conduits of external signals to generation of integrated cell and organ response (*1*). The primary transducers of activated GPCRs are heterotrimeric G proteins, comprised of distinct Gα and Gβγ subunits. While GPCRs are often classified according to the best coupled Gα subunit family, most GPCRs can couple, selectively, to members of multiple G protein subfamilies, and it is increasingly recognised that integrated signalling from multiple G proteins, and other transducers, plays an important role in governing complex cell responses (*2, 3*). However, the molecular basis for G protein selectivity of GPCRs remains poorly understood, with a major limitation being lack of structures of individual GPCRs bound to different G proteins.

Advances in single particle cryo-electron microscopy have enabled determination of agonist-activated GPCRs in complex with canonical transducer G proteins of the Gs, Gi/o and Gq/11 families using a range of biochemical approaches for stabilisation of these complexes (*4–8*). However, these have not yet translated to robust generation of complexes of GPCRs bound to more weakly coupled G proteins. A number of mechanisms, in addition to specific receptor-G protein interactions, have been proposed to contribute to G protein selectivity including the volume of the intracellular binding pocket in the receptor and the degree of conformational flexibility in TM6 (*9*). In particular, in most structures solved to date, the Gαs protein exhibits a bulkier C-terminal α5 helix (αH5) arising from a “hook” conformation of the far C-terminus that requires a larger binding pocket in the core of the receptor to be accommodated (*6, 10*). In these GPCR-Gs complex structures, TM6 is splayed further away from the core than is seen for Gi/o or Gq/11 complexes where these are the primary transducers. However, in more recent class A GPCR-Gs complexes greater divergence in the conformation of the intracellular TM helix ends has been observed (*11, 12*). Moreover, in the EP4 receptor, the C-terminal Gαs “hook” unwinds to enable novel engagement with the receptor (*11*) that could also enable binding of Gs to receptors that have narrower intracellular cavities when activated. The αH5 of Gi/o or Gq/11 proteins is less bulky and these proteins can be readily accommodated with smaller outward movement of TM6. It is clear that the nature of G protein engagement with GPCRs is complex and that individual receptor subfamilies can exhibit divergence in modes of G protein engagement, even for equivalent Gα proteins. This also raises questions on the mechanisms that contribute to G protein selectivity/promiscuity for individual GPCRs, which is critical for molecular understanding of biased agonist that can alter the pattern of G protein binding to GPCRs (*13*).

Recently, we solved structures of the glucagon receptor, a primarily Gs coupled receptor, in complex with Gs or Gi1 proteins, providing the first structural insight into G protein coupling pleiotropy (*14*). In contrast to expectation, the receptor backbone and intracellular pocket volume were equivalent regardless of the G protein bound, but with Gi2 binding within this cavity with fewer contacts. Currently, it is unclear how GPCRs, where Gq/11 proteins are the primary transducers, pleiotropically engage with Gs proteins.

The cholecystokinin (CCK) type 1 receptor (CCK1R) is a Gq/11 coupled class A GPCR localised on afferent vagal nerves that mediates the neuroendocrine peptide hormone actions of CCK on regulation of food intake and body weight (*15–17*). Through effects on gastric emptying and gut transit, in concert with stimulation of gall bladder contraction and pancreatic exocrine secretion, the CCK-CCK1R axis is a key physiologic servomechanism for optimal nutrient delivery and maintenance of body weight. CCK was also the first gut peptide shown to control satiety, and the CCK1R has been pursued as a potential target for treatment of obesity.

While, Gq/11 protein-dependent signalling has been the focus of pharmacological characterisation of the CCK1R, like most GPCRs, it is pleiotropically coupled and can initiate signalling via multiple transducers, including Gs- and G13-linked signaling, arrestin recruitment, and a wide array of other downstream effectors, including Ras, Raf, Rac, JNK, CDC42, p38, pERK, AKT, mTOR, S6 kinase, calcineurin, NFAT, and STATs (*9, 18, 19*). Mechanistic insight into the activation and transducer coupling of the CCK1R requires structural understanding of ligand binding and transducer engagement. However, no structures of the CCK1R, either in inactive or active states, have been solved. Moreover, we have recently demonstrated that increased cholesterol in the plasma membrane that routinely occurs in obese patients, impairs CCK-mediated Gq/11 protein signalling from the CCK1R (*20–22*). As such, obese patients are liable to be refractory to drugs that mimic CCK activation of this pathway and this has likely contributed to lack of clinical success of drugs developed using Gq/11 mediated pathways as the primary endpoint.

In this study, we used cryo-electron microscopy to determine structures of the CCK1R bound to the CCK peptide agonist, CCK-8 and two different transducer proteins; GαsGβ1γ2 and a chimeric Gq protein mimic (with Gβ1γ2) that was recently used to determine the structure of Gq/11 coupled 5HT_2A_ and orexin 2 (OX_2_) receptors (*23, 24*). As seen with other Gq/11-GPCR complexes, the Gq-α5 helix bound to a relatively narrow pocket in the CCK1R core. Surprisingly, the backbone of the CCK1R and volume of the G protein binding pocket was essentially equivalent when Gs was bound, with the Gs αH5 displaying a conformation that arises from “unwinding” of the far C-terminal residues, compared to canonically Gs coupled receptors. Thus, integrated changes in the conformations of both the receptor and G protein play critical roles in the promiscuous coupling of individual GPCRs.

## Results and Discussion

### CCK1R signalling is differentially modulated by the cellular environment

The CCK1R can couple to multiple G proteins including Gq/11 and Gs family proteins (*9*), but the potential importance of non-Gq/11 pathways is not clear. In HEK293s cells stably expressing the CCK1R, CCK-8 was ~1000-fold more potent in mobilisation of intracellular Ca^2+^ (iCa^2+^) downstream of Gq than Gs-mediated cAMP production (**Fig. 1A, 1I**), consistent with classification of CCK1R as a Gq/11 coupled receptor. As previously reported (*25*), increasing plasma membrane cholesterol (here delivered as a conjugate with MβCD) led to an ~10-fold loss of CCK-8 potency (**Fig. 1B, 1I**). In contrast, the increased cholesterol augmented peptide potency in cAMP production by ~10-fold (**Fig. 1B, 1I**) such that potency for the two pathways was only ~30-fold different. The increased potency for cAMP production was paralleled by a similar increase in whole cell binding affinity (**Fig. 1C, 1I**), which has previously been observed for CCK1R expressing cells in high cholesterol states (*25, 26*). Remarkably, in cells with genetic deletion of Gq/11 proteins there was also higher CCK-8 potency in cAMP production, where increasing cholesterol had no further effect (**Fig. 1D, 1I**). In these cells, however, the affinity of CCK-8 was similar to the parental cells, but cholesterol had reduced ability to increase binding affinity (**Fig. 1E, 1I**). In cells with genetic deletion of Gs, CCK-8 potency for iCa^2+^ mobilisation was similar to that seen with parental cells, as was the effect of cholesterol to reduce peptide potency (**Fig. 1F, 1I**). Interestingly, CCK-8 binding affinity was higher than seen with parental cells and was no longer sensitive to increased cholesterol (**Fig. 1G, 1I**). Collectively, these data illustrate that the CCK1R has a complex mode of G protein transducer engagement that is regulated by transducer expression levels and the local plasma membrane environment. Moreover, the data support the relevance of Gs coupling to the CCK1R with this also likely to contribute more to pathological signalling of the receptor. We next sought to understand the structural basis for pleiotropic coupling of CCK1R to Gq and Gs proteins.

**Fig. 1.**
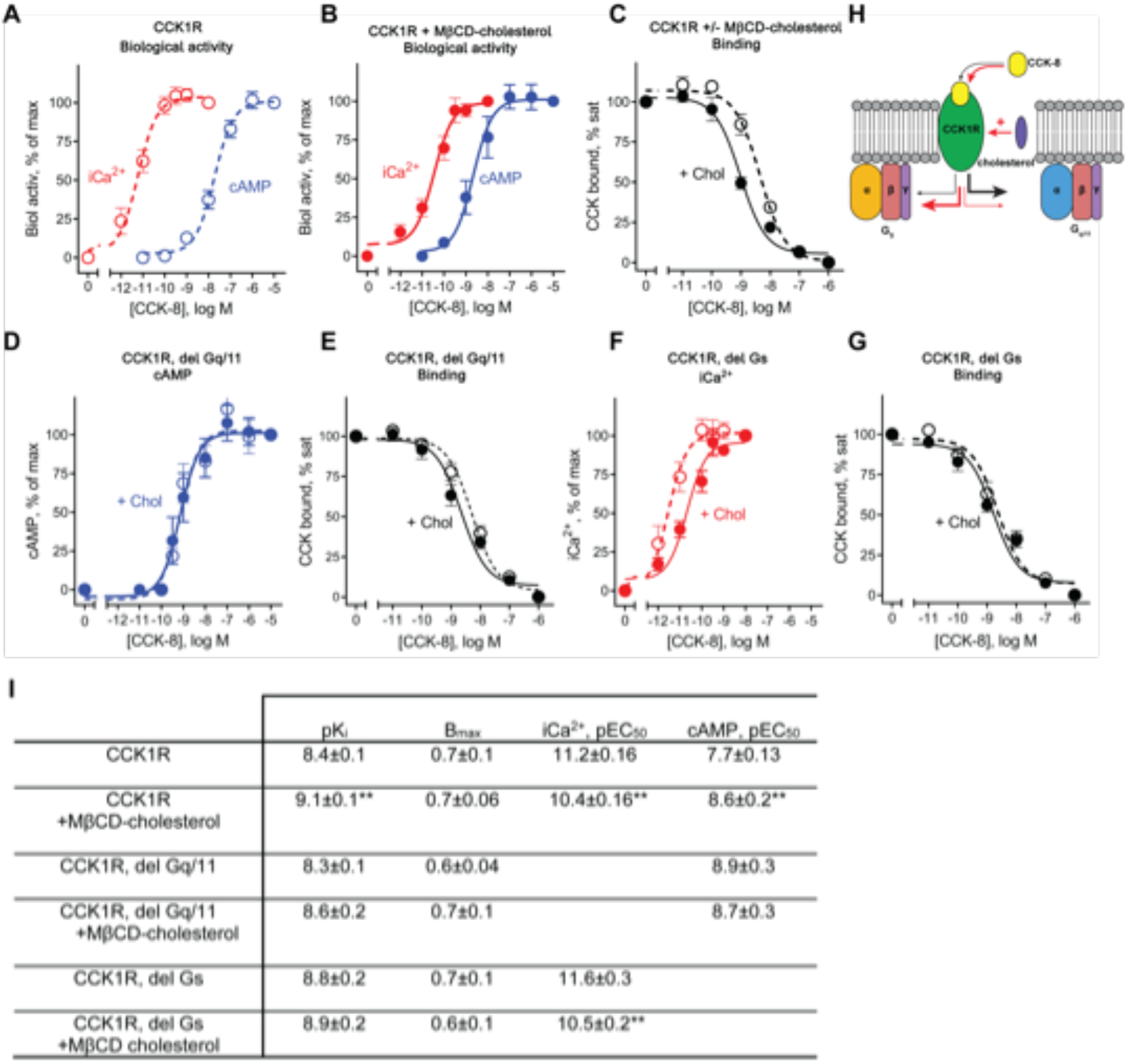
Effects of G protein association and cellular cholesterol on CCK-8 binding and biological activity. Shown are data for stable CCK1R-expressing HEK293s cell lines for parental cells or with deletion of Gq/11 or Gs proteins, in the absence or presence of increased cellular cholesterol by treatment with MβCD-cholesterol complex. Receptor density was not different between the cell lines. [**A**] In untreated cells, CCK was much more potent in stimulating iCa^2+^ mobilization than cAMP production. [**B**] After enhancing cellular cholesterol, CCK-8 potency in the iCa^2+^ mobilization assay was reduced but potency for cAMP production was increased. [**C**] Increased cellular cholesterol resulted in an increase in CCK-8 binding affinity. [**D**] In the cell line in which Gq/11 proteins were deleted, CCK-8 potency in the cAMP assay to CCK was increased and were not affected by cholesterol enhancement, whereas [**E**] CCK-8 binding affinity was not significantly different. [**F**] In the cell line in which Gs protein was deleted, CCK-8 potency in the iCa^2+^ assay and sensitivity to increasing cellular cholesterol were equivalent to parental cells, whereas CCK-8 binding affinity was insensitive to altered cholesterol. [**H**] Schematic of CCK1R signalling. Under conditions of normal membrane cholesterol CCK-8 signals predominantly via Gq/11 proteins with weak activation of Gs protein. With high cholesterol, there is increased Gs-mediated and decreased Gq/11-mediated signaling. [**I**] Quantitative pharmacology analysis of the data in panels A-G. Values are mean ± S.E.M. from 6-8 independent experiments performed in duplicate. **P<0.01; Significant differences were determined using a Mann-Whitney test for treated cells versus cells without MßCD-cholesterol treatment.

Binding of heterotrimeric G proteins to activated GPCRs is inherently unstable as the role of the GPCR is to act as a guanine nucleotide exchange factor (GEF) to rapidly activate and release the Gα and Gβ γprotein subunits, and prime subsequent second messenger activation events. As such, biochemical methods to stabilise binding of the Gα and Gβγ subunits to each other and to the activated receptors are required (*4–8*). The most robust approaches have been for GPCR complexes with Gs, where a combination of nanobody 35 (Nb35) with dominant negative Gαs, or mini-Gαs that lacks the mobile α-helical domain (AHD), has been used successfully for numerous receptors (*4, 10*). In contrast, complexes of GPCRs with Gq/11 proteins have been more difficult. The limited success has been with chimeric G proteins. In one approach, used with the muscarinic M_1_ receptor, chimeras of G11 with the αN of Gi were generated to enable the use of the short chain antibody, scFv16, to bridge this αN and the β-subunit of the obligate Gβγ dimer (*27*). For the 5HT2A and OX2 receptors, further engineering was required (*23, 24*). In this case a chimera of mini-Gs substituted with (i) Gq residues proximal to the receptor interface (including the C-terminal αH5) and (ii) the far αN of Gi (mGsqi; **Fig. S1I**).

In the current study, we used the mGsqi chimera in combination with scFv16 to maximise stability of ternary complexes with CCK-8 and CCK1R, where the mGsqi was fused to the receptor C-terminus, which was required for original complex formation. A 3C cleavage site was introduced prior to the G protein to allow for cleavage of the G protein following complex formation (**Fig. S1A**). For complexes with Gs, we utilised a dominant negative form of Gαs, with further stabilisation of the complex achieved using Nb35. The CCK1R for this complex was modified to include a HA-signal peptide, FLAG-epitope tag and the N-terminal sequence of the M4 mAChR that we have previously demonstrated to improve expression yields (*28, 29*). These sequences were followed by a 3C cleavage site to allow removal after affinity purification (**Fig. S1B**). The constructs (post cleavage equivalents) were not different from WT receptor in Gq-mediated iCa^2+^ mobilisation assays (**Fig. S1C**). Receptor, Gα (GαsDN; mGαsqI as a fusion with CCK1R) and Gβ1γ2 were co-expressed in Tni insect cells with complex formation initiated by the addition of 10 μM CCK-8. Complexes were further stabilized through addition of apyrase, to remove guanine nucleotides, and by addition of either scFv16 (mGsqi) or Nb35 (GsDN). Complexes were solubilised in LMNG/CHS, followed by anti-FLAG affinity purification, treatment with 3C enzyme and separation by size exclusion chromatography (SEC) to yield monodisperse peaks containing the protein complex (**Fig. S1D–S1F and S1G–S1H**). Following vitrification, samples were imaged on a Titan Krios with data processed to yield 3D consensus reconstructions of 2.5 Å and 2.0 Å resolution at gold standard FSC 0.143, respectively, for the CCK-8/CCK1R/mGsqi and CCK-8/CCK1R/GsDN complexes (**Fig. S1J, 1K; Fig. 2A – 2F**).

**Fig. 2.**
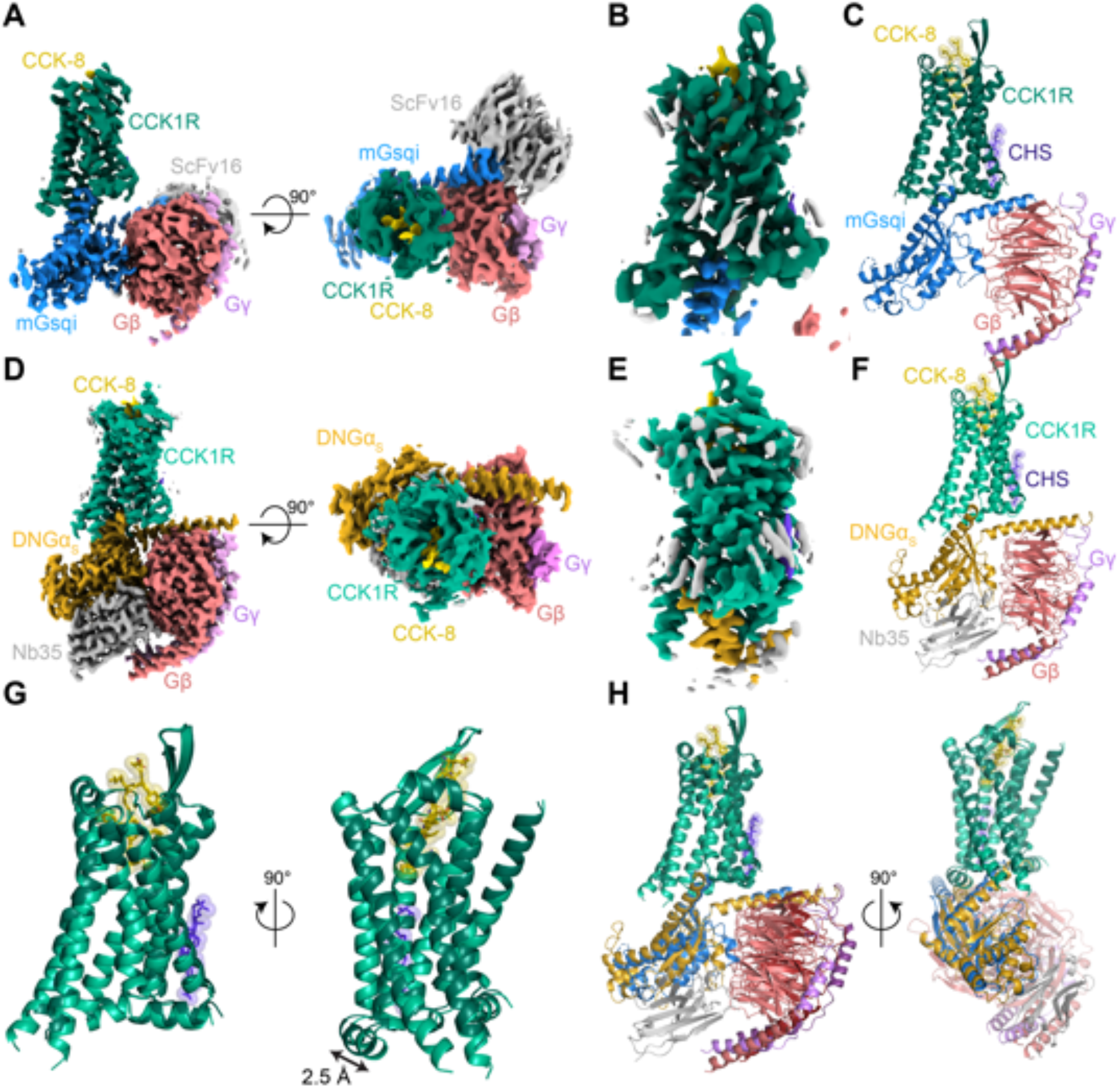
Cryo-EM structure of CCK-8/CCK1R in complex with DNGαs/Gβ1γ2/Nb35 or mGαsqi/Gβ1γ2/scFv16. [**A**] Consensus cryo-EM map of the CCK-8/CCK1R/mGαsqi/ Gβ1γ2/scFv16 complex resolved to 2.45 Å (FSC 0.143). [**B**] Cryo-EM map following focused refinement of the receptor resolved to 2.5 Å (FSC 0.143). [**C**] Molecular model of the complex (the scFv16 was omitted from modelling). [**D**] Consensus cryo-EM map of the CCK-8/CCK1R/DNGαs/Gβ1γ2/Nb35 complex resolved to 1.95 Å (FSC 0.143). [**E**] Cryo-EM map following focused refinement of the receptor resolved to 2.1 Å (FSC 0.143). [**C**] Molecular model of the complex. [**G**], [**H**] Alignment of the two structures. [**G**] Alignment of receptor and CCK-8 peptide. The largest difference in the CCK1R was in the location of ICL2 that is further away from the receptor core in the complex with the Gq-mimetic protein. [**H**] Alignment of the full complex illustrating differences in the engagement and orientation of the G proteins. The maps and models are coloured according to the labels on the figure. The receptor and G proteins are displayed in ribbon format. The CCK-8 peptide and modelled cholesterol are displayed in ball and stick representation.

For the complex with the Gq-mimetic, local resolution was highest for Gβ and Gα subunits that are stabilised by scFv16 with lowest resolution for the extracellular face of the receptor and peptide (**Fig. S2A**). Additional focused refinements were performed on the receptor and G protein (**Fig. S2B, S2C**) leading to substantially improved resolution of these domains that allowed modelling of the Ras domain of the Gα, Gβ, Gγ, CCK1R and CCK-8, including side chain rotamers for the receptor with the exception of ICL3 residues 244 to 301 that were poorly resolved and not modelled (**Fig. 2C, 2G, 2H**; **Fig. S2D**).

For the complex with Gs, local resolution was highest for the G protein and G protein-receptor interface, with lowest resolution at the extracellular face of the receptor (**Fig. S2E**). Additional focused refinement of the receptor provided improvement to the local resolution, including the peptide binding site (**Fig. S2F**), allowing accurate modelling of most side chain rotamers and also waters within the binding pocket and receptor-G protein interface (**Fig. 2F–2H; Fig. 3A; Fig. S2G**).

**Fig. 3.**
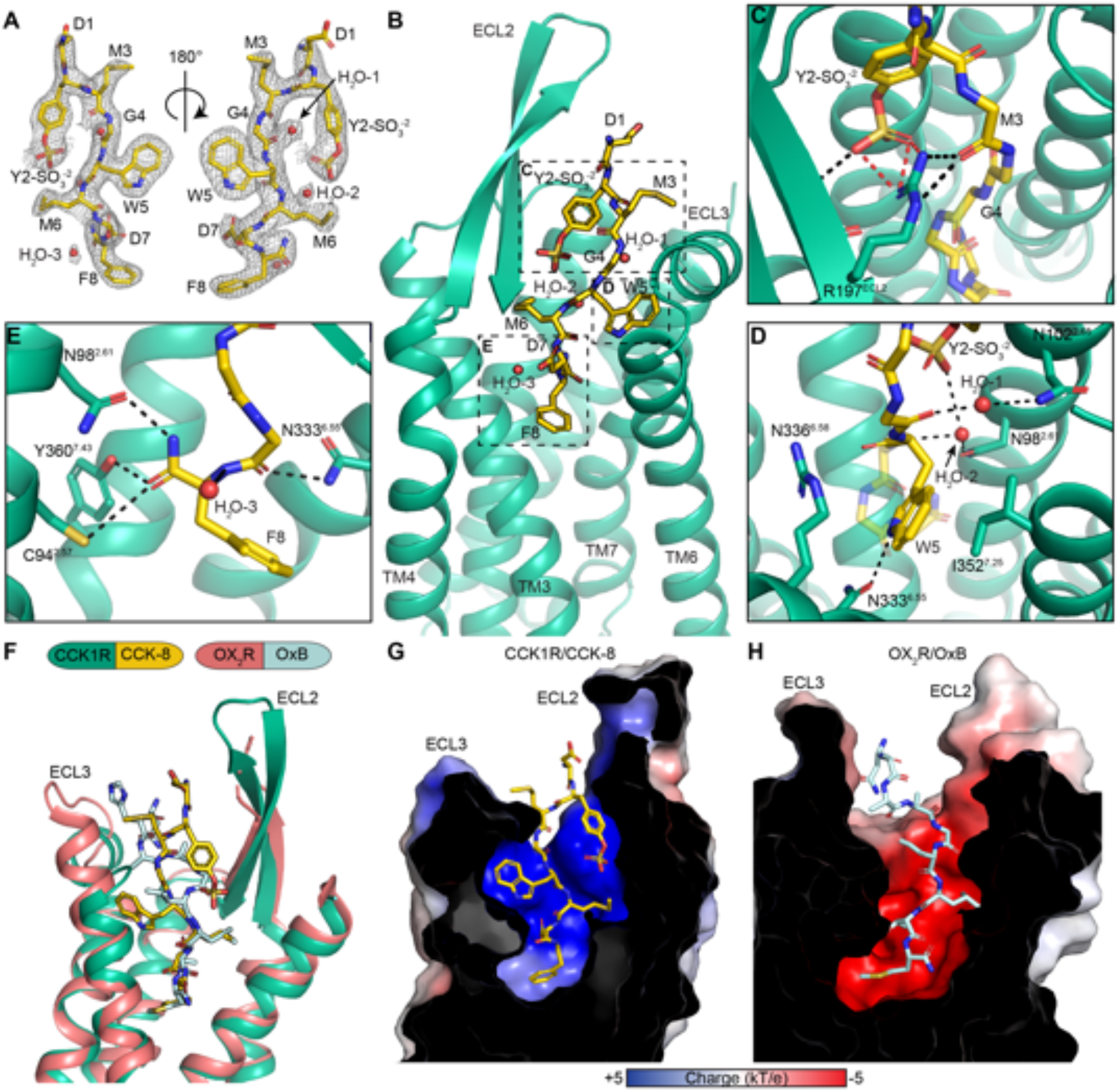
CCK1R interactions with CCK-8. [**A**] EM density for the CCK-8 peptide ligand (yellow, stick representation coloured by heteroatom) and proximal waters (red spheres) zoned at 1.8 Å. [**B**] CCK-8 (yellow) is bound with the C-terminus of the peptide buried within the TM bundle and makes extensive interactions with CCK1R (green, ribbon format). [**C**] Y2-SO_3_^CCK^ makes interactions with R197^ECL2^ and C196^ECL2^. [**D**] W5^CCK^ makes interactions with N333^6.55^ and I352^7.25^. **[E]** The terminal F8-NH2CCK forms hydrogen bonds with C94^2.57^, N98^2.61^, L356^7.39^ ^61^ and Y360^7.43^. **[F]** Comparison of the CCK-8/CCK1R with OxB/OX_2_R (blue ligand, pink receptor) binding pockets. The peptides overlap in the location of the amidated C-terminal tetrapeptide but differ in conformation and position of N-terminal amino acids. CCK-8 interacts more with ECL2 than ECL3 while the reverse is true for OxB. [**G**] Surface electrostatic potential of the CCK1R binding pocket. CCK1R has a very positive surface charge. [**H**] In contrast the OX_2_R binding pocket has a very negative surface potential.

The backbone of the receptor and location of CCK-8 in the binding pocket were highly similar for both the Gq-mimetic and Gs bound complexes with root mean square deviations of 0.6 Å for the receptor and 0.3 Å for the peptide. The greatest divergence between structures was observed in the position of ICL2 that was translated 2.5 Å away from the receptor core in the complex with the Gq-mimetic (**Fig. 2G; Video S1**). While ECL1 was unstructured, ECL2 formed a twisted β-hairpin and ECL3 a short α-helix. On the intracellular face, ICL2 presented as a short α-helix (**Fig. 2G**). While both G proteins penetrated the core of the receptor to a similar depth, there were translational and rotational differences in the orientation of G proteins relative to the receptor (**Fig. 2H**), described in detail below.

### CCK-8 binding

CCK-8 bound into the consensus structures of both CCK1R-DNGs and CCK1R-mGsqi in essentially identical poses (**Fig. S3**). As the binding site resolution was higher with the Gs-complex, the interactions between CCK-8 and the receptor are described for this structure (**Fig. 3, Table 1**). The CCK-8 peptide is bound to the receptor in an extended conformation, with the C-terminus buried deep within the core of CCK1R and the N-terminus pointing up and out of the ligand binding cavity (**Fig. 3B**). The amidated C-terminal F8^CCK^ forms a hydrogen bond network to C94^2.57^, N98^2.61^ and Y360^7.43^ of the CCK1R, as well as van der Waals interactions with L356^7.39^ (**Fig. 3E, Table 1**). The C-terminus is further stabilised by the side chain of D7^CCK^, which forms salt bridges to H210^5.39^ and R336^6.58^ as well as a hydrogen bond to Y176^4.60^. The centre of the peptide is primarily coordinated by a series of van der Waals interactions, of note W5^CCK^ forms a hydrogen bond to N333^6.55^ as well as van der Waals interactions to I352^7.25^ (**Fig. 3D, Table 1**). The sulphated tyrosine (Y2-SO_3_^CCK^), although close to the N-terminus of the peptide, is folded over pointing back down towards the core of the receptor. It forms a number of salt bridges to R197^ECL2^ of the β-strand of ECL2, as well as a hydrogen bond to the backbone of C196 ^ECL2^. R197^ECL2^ is further coordinated by hydrogen bonds to the backbone of M6^CCK^ of CCK-8 (**Fig. 3C**). The high resolution of this structure enabled modelling of three water molecules within the CCK-8 binding pocket, one of which (H_2_O-2) is coordinated by the SO_3_ group of Y2-SO_3_^CCK^, and also interacts with the backbone of M6^CCK^ and N98^2.61^ of CCK1R. The location of CCK-8, which is deep within the TM binding pocket, is observed in other class A peptide-bound receptor structures, such as the OX_2_R, which also binds a C-terminally amidated peptide (OxB) (*24*). The similar depth of OxB within the OX_2_R binding pocket also enables hydrogen bonding to Y^7.43^. OxB also extends out towards the extracellular surface, yet orients towards ECL3 at the N-terminus, whereas CCK-8 forms interactions with ECL2 of CCK1R (**Fig. 3F**). The negatively charged nature of CCK-8 side chains favours the relatively positively charged pocket of CCK1R as opposed to the negatively charged pocket observed for OX_2_R (**Fig. 3G–H**).

**Table 1.** List of contacts between CCK-8 and CCK1R.

| Non-bonded contacts |  |  | Non-bonded contacts continued |  |  |
| --- | --- | --- | --- | --- | --- |
| CCK-8 | CCK1R | Distance (Å) | CCK-8 | CCK1R | Distance (Å) |
| D1 | F185<sup>ECL2</sup> | 3.69 | D7 | F198<sup>ECL2</sup> | 3.39 |
| D1 | M195<sup>ECL2</sup> | 3.50 | D7 | HIS210<sup>5.39</sup> | 3.36 |
| Y2-SO<sub>3</sub> | P101<sup>2.64</sup> | 3.59 | F8 | L356<sup>7.39</sup> | 3.50 |
| Y2-SO<sub>3</sub> | K105<sup>2.68</sup> | 3.33 | F8-NH<sub>2</sub> | N98<sup>2.61</sup> | 2.8 |
| Y2-SO<sub>3</sub> | D106<sup>ECL1</sup> | 3.61 | **Hydrogen Bonds** |  |  |
| Y2-SO<sub>3</sub> | M195<sup>ECL2</sup> | 3.53 | **CCK-8** | **CCK1R** | **Distance (Å)** |
| M3 | M195<sup>ECL2</sup> | 3.47 | Y2-SO<sub>3</sub> | C196<sup>ECL2</sup> | 2.94 |
| M3 | E344<sup>7.29</sup> | 3.35 | M3 | R197<sup>ECL2</sup> | 2.96 |
| G4 | R197<sup>ECL2</sup> | 3.22 | G4 | S348<sup>7.31</sup> | 2.94 |
| G4 | E344<sup>7.29</sup> | 3.81 | W5 | N333<sup>6.55</sup> | 3.30 |
| W5 | R197<sup>ECL2</sup> | 3.46 | D7 | Y176<sup>4.60</sup> | 3.35 |
| W5 | R197<sup>ECL2</sup> | 3.43 | D7 | N333<sup>6.55</sup> | 2.94 |
| W5 | R336<sup>6.58</sup> | 3.54 | F8 | C94<sup>2.57</sup> | 3.80 |
| W5 | L347<sup>7.30</sup> | 3.61 | F8 | Y360<sup>7.43</sup> | 2.50 |
| W5 | I352<sup>7.35</sup> | 3.42 | **Salt bridges** |  |  |
| M6 | F97<sup>2.60</sup> | 3.26 | **CCK-8** | **CCK1R** | **Distance (Å)** |
| M6 | N98<sup>2.61</sup> | 3.60 | Y2-SO<sub>3</sub> | R197<sup>ECL2</sup> | 2.60 |
| M6 | T118<sup>3.29</sup> | 3.77 | Y2-SO<sub>3</sub> | R197<sup>ECL2</sup> | 2.74 |
| M6 | M121<sup>3.32</sup> | 3.75 | D7 | R336<sup>6.58</sup> | 3.60 |
| M6 | C196<sup>ECL2</sup> | 3.36 |  |  |  |

CCK-8 interactions are, in part, supported by previous mutagenesis studies. Select mutants of C94^2.57^, R197^ECL2^, N333^6.55^, I352^7.25^, L356^7.39^, and Y360^7.43^ cause a decrease in the potency of CCK peptides that range from 30-fold to 9,300-fold (*30–32*). However, it is important to note that the binding mode of CCK-8 to its receptor has been controversial with the peptide C-terminus predicted to occupy either a shallow or deep pose, depending upon the method of investigation (*31, 33*). We have recently speculated that the deep pose might be equivalent to the higher affinity state seen in the presence of high cholesterol. The peptide location in the consensus maps is consistent with the deeper pose. In the absence of Gq/11 proteins, CCK-8 potency for Gs-mediated cAMP production is higher and insensitive to increased cholesterol. As such, it is possible that CCK-8 might favour the more stable, deeper pose when bound to Gs. For both preparations, the complexes were solubilised in detergent supplemented with CHS and consequently may more closely mimic a high cholesterol type environment leading to enrichment of the deeper binding mode. The CCK1R contains 4 consensus cholesterol binding motifs (*25*) and both complexes are surrounded by annular lipids (**Fig. S4D, S4E**), including density that could be modelled with CHS located at the interface of TM2 and TM4 (**Fig. S4A–S4C**). The functionally relevant allosteric cholesterol binding domain has been localised to a CRAC motif low in TM3 (*25, 34*). In the consensus structures, weaker densities that might correspond to cholesterol are present in the cryo-EM maps, with the density extending to ICL2 (**Fig. S4D, S4E**), suggesting that bound cholesterol may modulate the conformational dynamics of this loop. In other class A GPCRs, including the β_2_-AR (*35*), hydrophobic residues of ICL2 interact with the junction of the αN and αH5 and have been linked to G protein activation. The density for ICL2 is well resolved for the Gs-complex, but there is additional density in the Gq-bound complex that might indicate higher relative dynamics (**Fig. S4F**), and this was also observed in 3D multivariance analysis (3DVA) of the principal components of motion within the cryo-EM data (**Video S2**). In both CCK1R complexes, the receptor was highly dynamic, undergoing twisting and rocking motions, and this was particularly true for the extracellular regions of the receptor and CCK-8 peptide binding site (**Video S2**). This likely reflects differential stability of the peptide in the deep binding pose, even when stabilised by G protein binding. Nonetheless, additional work will be required to better understand the structural basis for the differences reported in modes of peptide binding to the CCK1R and how this is regulated by different lipids.

### CCK1R activation mechanism

The transmembrane core of the active-state CCK1R is similar to other active-state class A GPCR structures (*6*) with the intracellular side of TM6 occupying a similar location to other active, Gq/11-coupled, receptors (*23, 24, 27*), creating a binding site for the insertion of the α5 helix of the G proteins. While there are no inactive-state structures available for the CCK1R, the rotameric positions of residues in conserved class A activation motifs such as the CWxP, PI(T)F, NPxxY, and E/DRY motifs exhibited strong overlap with the equivalent residues in the active OX_2_R complex (*24*) (**Fig. S5**). As such, the transitions observed between the inactive and active OX_2_R serve as a template for the likely reorganisation of residues in these activation motifs. At the bottom of the peptide binding pocket the aromatic edge of F8^CCK^ interacts with F330^6.52^ that in turn interacts with W326^6.48^ of the CWxP motif (**Fig. S5A**). The position of W326^6.48^ is such that F322^6.44^ of the PI(T)F motif moves outward. This outward rotation of F322^6.44^ is consistent with other class A GPCRs and is proposed to initiate the outward movement of TM6 that is necessary for G protein binding (**Fig. S5B**). Further below the PI(T)F motif is residue Y370^7.53^ within the NPxxY motif, which moves inwards, promoting an interaction with Y229^5.58^ through a bridging water molecule (**Fig. S5C**). This tyrosine “water lock” is speculated to stabilize the active conformation of class A GPCRs (*37*). The stabilisation of Y229^5.58^ by the “water-lock” allows this residue to interact with R139^3.50^ from the E/DRY motif, following release from an ionic interaction proposed to occur between R139^3.50^ and E138^3.49^ in inactive state structures (**Fig. S5D**).

### CCK1R-G protein interface

Overall, the structures of CCK1R/Gs and CCK1R/mGsqi are similar and display the prototypical receptor/G protein interface, however they also exhibit marked differences in their mode of G protein binding. When the structures are overlayed on the receptor, the G proteins display substantive differences in their orientation, with differences in the Gα protein propagated to the Gβγ subunits (**Fig. 2H, Fig. 4A; Fig. S6A, 6B; Video S1**). For example, while the base of the Gα α5 helices overlay, the C-terminus of this helix in Gs is rotated ~9.5° outwards from the core of the receptor, relative to mGsqi (**Fig. 4C, 4D; Video S1**). In combination with unwinding of the C-terminal “hook” of Gs (discussed below), the C-terminal residues are oriented out from the receptor core between TM6 and TM7/H8 and this allows the conserved L393^H5.25^ (superscript, CGN G protein numbering system (*38*); **Fig. 4B**) to form van der Waals interactions with M373^7.56^ and R310^6.32^ (**Fig. 4E; Table 2**). The C-terminus of the α5 helix of Gs is further stabilised by a hydrogen bond from N374^6.47^ to the backbone of Q390^H5.22^. Furthermore, the conserved L388^H5.20^ sits in between I143^3.54^ and K308^6.30^, forming close contacts and potentially contributing to stabilisation of TM3 and TM6 (**Fig. 4E; Table 2**). The rotation inwards of the α5 helix of mGsqi places the conserved L393^H5.25^, much further inwards (7.6 Å), allowing it to sit in a hydrophobic pocket formed by V311^6.33^ and L315^6.37^, and in close proximity to R139^3.50^ of the DRY motif (**Fig. 4F; Table 2**). This shift inwards results in fewer interactions with TM7 and H8, compared to Gαs, with only R376^8.49^ forming a hydrogen bond to R389^H5.21^. This shift inwards allows the conserved Y391^H5.23^ to hydrogen bond to Q153^ICL2^, potentially contributing to conformational positioning of ICL2 (**Fig. 4F; Table 2**); this bond is not observed in the Gs-bound structure. The non-conserved N387^H5.19^ forms a hydrogen bond to the backbone of A142^3.53^ at the base of TM3 (**Fig. 4F; Table 2**). In Gs this corresponds to H387^H5.19^, which can also interact with A142^3.53^ but potentially forms an additional hydrogen bond to R150^ICL2^ (3.4 Å), further contributing to stabilisation of ICL2. In 3DVA of the cryo-EM data to extract principal components of motion for each of the CCK1R complexes, the α5 helix C-terminus was more stable for the mGsqi protein than the corresponding region of the Gs protein (**Video S3**), despite greater motion overall, when comparing the entire G protein relative to the receptor (**Video S2-S4**).

**Table 2.** List of contacts between CCK1R and G proteins.

**CCK1R/Gαs contacts**
| Non-bonded contacts |  |  | Non-bonded contacts continued |  |  |
| --- | --- | --- | --- | --- | --- |
| Gas | CCK1R | Distance (Å) | Gas | CCK1R | Distance (Å) |
| R38 | R150<sup>ICL2</sup> | 3.07 | Q390 | N374<sup>8.47</sup> | 2.93 |
| H41 | L147<sup>ICL2</sup> | 3.61 | Q390 | T76<sup>2.39</sup> | 3.72 |
| H41 | R150<sup>ICL2</sup> | 3.58 | Y391 | V311<sup>6.33</sup> | 3.82 |
| D215 | Q148<sup>ICL2</sup> | 2.99 | Y391 | R139<sup>3.50</sup> | 3.66 |
| V217 | Q148<sup>ICL2</sup> | 3.34 | L393 | M373<sup>7.56</sup> | 3.22 |
| D354 | A302<sup>6.24</sup> | 3.59 | L393 | R310<sup>6.32</sup> | 3.69 |
| Y358 | N304<sup>6.26</sup> | 3.80 | L394 | K375<sup>8.48</sup> | 3.34 |
| F376 | L147<sup>ICL2</sup> | 3.56 | L394 | R378<sup>8.51</sup> | 3.01 |
| R380 | L147<sup>ICL2</sup> | 3.85 | R389 | R376<sup>8.51</sup> | 3.48 |
| R380 | P146<sup>ICL2</sup> | 3.89 | **Hydrogen Bonds** |  |  |
| R380 | E235<sup>8.48</sup> | 3.37 | **Gas** | **CCK1R** | **Distance (Å)** |
| R380 | C144<sup>3.55</sup> | 2.63 | R38 | R150<sup>ICL2</sup> | 3.07 |
| I383 | L147<sup>ICL2</sup> | 3.66 | R38 | R150<sup>ICL2</sup> | 3.29 |
| Q384 | P146<sup>ICL2</sup> | 3.68 | D215 | Q148<sup>ICL2</sup> | 2.99 |
| Q384 | I143<sup>3.54</sup> | 2.93 | R380 | C144<sup>3.55</sup> | 2.63 |
| Q384 | E235<sup>5.64</sup> | 3.63 | H387 | A142<sup>3.53</sup> | 2.75 |
| R385 | N304<sup>6.26</sup> | 3.82 | Q390 | N374<sup>8.47</sup> | 3.34 |
| H387 | A142<sup>3.53</sup> | 2.75 | **Salt bridges** |  |  |
| H387 | P146<sup>ICL2</sup> | 3.58 | **Gas** | **CCK1R** | **Distance (Å)** |
| H387 | R150<sup>ICL2</sup> | 3.40 | R380 | E235<sup>5.64</sup> | 3.37 |
| L388 | I143<sup>3.54</sup> | 3.77 |  |  |  |
| L388 | K308<sup>6.30</sup> | 3.66 |  |  |  |

**CCK1R/mGsqi contacts**
| Non-bonded contacts |  |  | Non-bonded contacts continued |  |  |
| --- | --- | --- | --- | --- | --- |
| mGsqi | CCK1R | Distance (Å) | mGsqi | CCK1R | Distance (Å) |
| R38 | R150<sup>ICL2</sup> | 3.69 | Y391 | R139<sup>3.50</sup> | 3.68 |
| R38 | T154<sup>4.38</sup> | 3.57 | Y391 | A142<sup>3.53</sup> | 3.63 |
| R38 | R150<sup>ICL2</sup> | 3.27 | Y391 | Q153<sup>ICL2</sup> | 2.34 |
| R38 | V151<sup>ICL2</sup> | 2.98 | N392 | N374<sup>8.47</sup> | 3.32 |
| R38 | T154<sup>4.38</sup> | 3.48 | N392 | T76<sup>2.39</sup> | 3.49 |
| R39 | V151<sup>ICL2</sup> | 3.38 | L393 | V311<sup>6.33</sup> | 3.25 |
| L41 | L147<sup>ICL2</sup> | 3.67 | L393 | R139<sup>3.50</sup> | 3.42 |
| L41 | R150<sup>ICL2</sup> | 3.41 | L393 | L315<sup>6.37</sup> | 3.79 |
| V217 | L147<sup>ICL2</sup> | 3.75 | **Hydrogen bonds contacts** |  |  |
| K380 | L147<sup>ICL2</sup> | 3.69 | **mGsqi** | **CCK1R** | **Distance (Å)** |
| I383 | P146<sup>ICL2</sup> | 3.62 | R38 | R150<sup>ICL2</sup> | 3.27 |
| I383 | L147<sup>ICL2</sup> | 3.74 | R38 | V151<sup>ICL2</sup> | 2.98 |
| I383 | R150<sup>ICL2</sup> | 3.64 | N387 | A142<sup>3.53</sup> | 3.38 |
| L384 | I143<sup>3.54</sup> | 3.26 | R389 | R376<sup>8.49</sup> | 3.57 |
| N387 | A142<sup>3.53</sup> | 3.72 | Y391 | Q153<sup>ICL2</sup> | 2.34 |
| R389 | R376<sup>8.49</sup> | 3.57 |  |  |  |
| E390 | T76<sup>2.39</sup> | 3.38 |  |  |  |

**Fig. 4.**
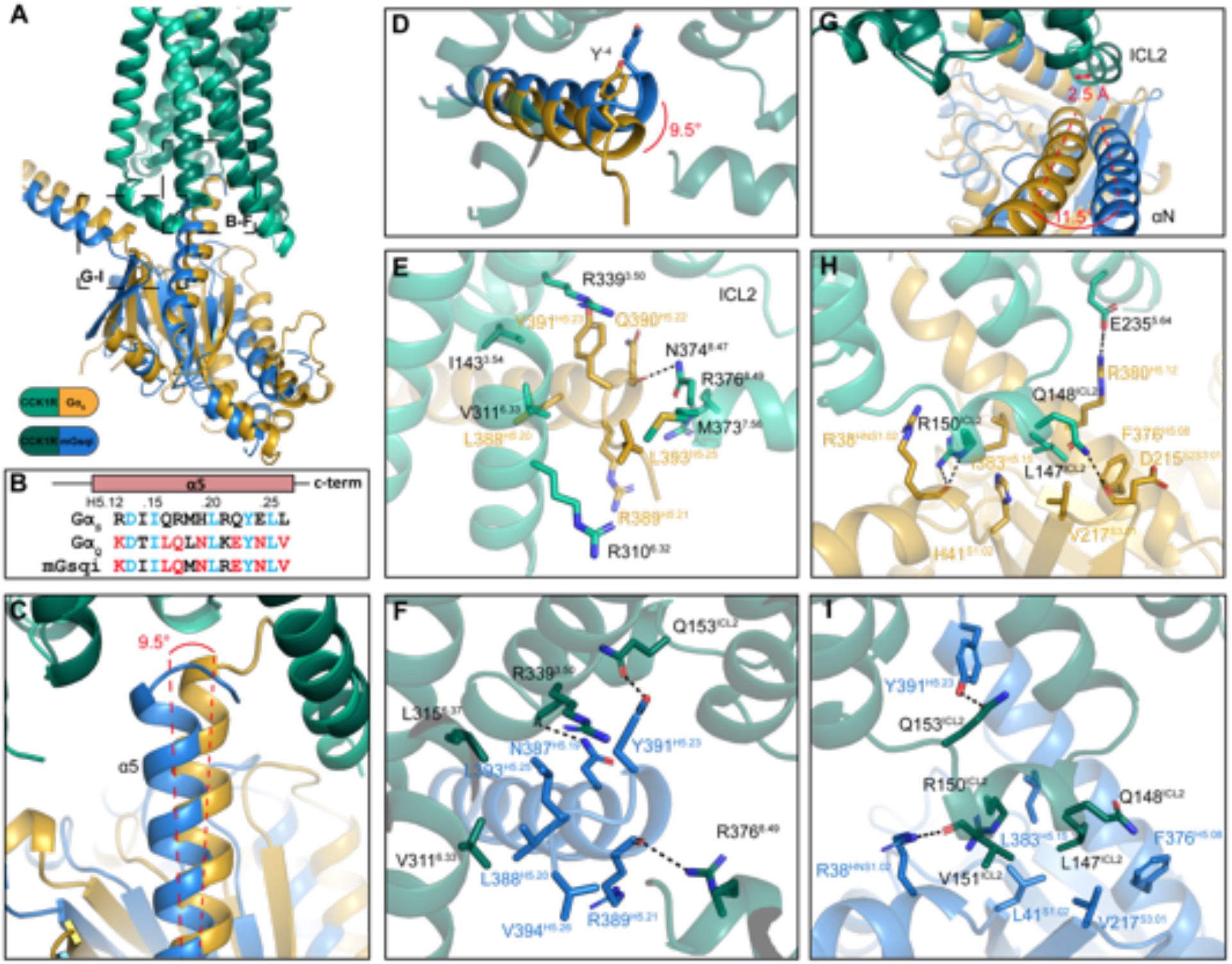
G protein interactions with activated CCK1R. [**A**] Overlay of Gαsubunits bound to the CCK1R highlighting the position of the magnified section in panels B-G. [B] the sequences of the GαC-terminus (α5 helix) for Gαs, Gαq or the chimeric mGαsqi protein. Residues common to both parental G proteins are coloured light blue, amino acids unique to Gs are coloured black and those of Gq are coloured red. [**C**], [**D**] There is an ~9.5-degree difference in the position of the C-terminus of the two G proteins relative to the origin of the α5 helix. [**E, F, H, I**] Gαs protein [**E, H**] and the Gαq-mimetic protein [**F, I**] form distinct interactions with CCK1R, illustrated from the top of the α5 helix [**E, F**] or the junction of the αN and α5 helices that interacts with ICL2 of the CCK1R [**H, I**]. Protein backbone is illustrated in ribbon format with Gαs in gold (CCK1R, light green) and mGαsqi in blue (CCK1R, dark green). Side chains from the G protein or receptor that interact are displayed in stick format, coloured by heteroatom. Dashed lines indicate H-bonds. G protein residues are numbered according to the CGN G protein numbering system (*38*).

Major differences were also observed between the consensus structures in the location and orientation of the αN helix. For mGsqi, the αN helix is rotated outwards from the receptor by ~11.5° (**Fig. 4G**) that is accompanied by a 2.5 Å shift in ICL2 away from the receptor TM bundle (**Fig. 2G, Fig. 4G; Video S1**). In both structures L147^ICL2^ is buried in a hydrophobic pocket within the α subunit, formed by the α5 helix, β2-β3 loop and the top of the β1 strand (**Fig. 4H, 4I**). For Gs L147^ICL2^ is buried deeper into the pocket, interacting with H41^S1.02^, V217^S3.01^, F376^H5.08^ and I383^H5.15^. Moreover, in the Gs-bound complex, ICL2 is further stabilised by hydrogen bonds between Q148^ICL2^ and R150^ICL2^ of the receptor and D215^S2S3.01^ and R38^HNS1.02^ of the G protein, respectively (**Fig. 4H**). For mGsqi, L147^ICL2^ also forms interactions within this hydrophobic pocket, which is comprised of L41^S1.02^, V217^S3.01^, F376^H5.08^ and L383^H5.15^, yet does not bind as deeply (**Fig. 4I**). ICL2 in this complex is further stabilised by a hydrogen bond between R38 ^HNS1.02^ and the backbone of V151^ICL2^ and the previously described hydrogen bond between Q153^ICL2^ and Y391^H5.23^ (**Fig. 4I**).

In the 3DVA, the mGsqi protein heterotrimer (with the exception of the more stable α5 helix) undergoes substantially greater movements than seen for the Gs-bound complex (**Video S4**). The more limited motion of Gs is likely due to transient interactions between ICL3 and the S6-H4 loop of the Gα-subunit that is relatively stable in most of the principal components of motion in the 3DVA (**Video S2**). Although not well resolved in either structure, there is more robust density for the ends of TM5 and TM6 in the Gs-bound complex versus the mGsqi-bound complex in the consensus reconstructions (**Fig. 2**), consistent with the observed interaction.

For both complexes, there is only limited contact with the Gβ1 subunit with only poor density in the cryo-EM maps for sidechains of residues located at the receptor-Gβ interface.

### Comparison of CCK1R G protein engagement with coupling of canonical G protein partners in other GPCRs

#### Gq/11 family proteins

The CCK1R is preferentially coupled to Gq/11 proteins. Only 3 other structures of GPCRs (5HT_2A_R, OX_2_R, M1 AChR) bound to Gq-mimetic proteins have been determined (*23, 24, 27*), where Gq or G11is also the primary transducer. In each of these receptors, TM6 remains in an extended helical conformation that is translated out at the intracellular base relative to inactive structures (*23, 24*); the G protein α5 helix binds into the narrow cavity formed by this translational movement (**Fig. S5, S7**). Overlay of the CCK1R/mGsqi complex structure with each of these other receptor-G protein complexes revealed good correlation in the location and orientation of the G protein, and in the conformation of the far C-terminal residues of the α5 helix (**Fig. S7A–S7C**). The largest difference was observed between the CCK1R/mGqsi and the M1 AChR/G11/i complexes, where there both the α5 and αN helices were translated relative to each other (**Fig. S7C**). The M1 AChR/G11/i is the only GPCR-Gq/11 complex solved where the Gα subunit is primarily the Gq/11 sequence, and this could contribute to the observed differences. However, it is important to note that in the 3DVA of the CCK1R/mGsqi cryo-EM data all the conformations presented in the available consensus structures of the active M1 AChR, OX_2_R and 5HT_2A_R are sampled (**Video S2, S4**). As such, sample preparation and vitrification conditions could also contribute to the observed differences, in addition to the potential for real distinctions in the most stable conformations for each complex.

#### Gs proteins

The Gs protein heterotrimer was the first transducer to be solved in complex with an activated GPCR (*39*), and multiple complexes with Gs have now been determined for class A and class B GPCRs where Gs is the primary transducer (*7, 10–12, 40–44*). These structures have revealed both common and diverse features that govern Gs binding. In the inactive GDP-bound Gαs protein, the α5 helix interacts with the αN helix and β1 sheet, with the far C-terminus assuming the bulky “hook” conformation that is prototypical of the active G protein conformation (**Fig. 5E, 5F**). In complex with activated GPCRs, the α5 helix is rotated anti-clockwise and translated away from the nucleotide binding site (**Video S5**). In most class A GPCR complexes, and all class B GPCR complexes, the bulky “hook” conformation is maintained with binding into the receptor intracellular binding cavity, primarily enabled by a large outward movement at the base of TM6 following receptor activation (*7, 10*; **Fig. 5**). In prototypical receptors, such as the β-ARs or the adenosine A_2A_R, this is facilitated by kinking of TM6 (**Fig. 5A, 5G, 5I**), and this is even more marked in class B GPCRs (*40*; **Fig. 5K, 5L**). This was initially thought to be a required feature for Gs coupling, at least for receptors where Gs is the primary transducer (*7, 9, 10*). Recent structures of class A GPCRs from different evolutionary subclades have demonstrated that these receptors can bind and activate Gs through alternative mechanisms. In both the bile acid receptor, GPBAR (*12*), and the prostanoid receptor, EP4R (*11*), TM6 is an unkinked extended α-helix, similar to the TM6 conformation in structures of the active Gq/11 coupled receptors (**Fig. 5H, 5J; Fig. S7**), with a correspondingly narrow pocket for binding of the C-terminal α5 helix. In the GPBAR-Gs complex, the α5 helix is in the prototypic “hook” conformation and this is accommodated by a shallower engagement with the receptor core that is nonetheless stabilised by more extensive interactions of the Ras domain with ICL3 and TM5 that has a markedly extended, stable, helix (*12*); a similar elongated TM5 helix and extended interactions with the Ras domain is observed in the GPR52-Gs structure (*41*). Nonetheless, the bulky “hook” conformation is maintained in these structures, suggesting that it is energetically favoured for Gs proteins. In contrast, in the CCK1R-Gs structure, the far C-terminal residues are “unwound” leading to a less bulky conformation and the α5 helix binds to the same depth as both the mGsqi protein in the CCK1R structure and the Gs protein in prototypical structures, with the C-terminal hydrophobic Leu residues oriented towards the lipid/detergent interface (**Fig. 5A, 5B, 5D**). Intriguingly, in the EP4R that preferentially couples to Gs, unwinding of the “hook” also occurs (**Video S5**), enabling the α5 helix to bind to similar depth in the narrower receptor cavity, although here the C-terminal residues orient between the base of TM7 and TM1 (*11*; **Fig. 5J**). These data demonstrate that, while there is a common mode for Gs to engage with GPCRs, functional engagement can occur by multiple divergent mechanisms in a receptor-dependent manner for both canonically coupled GPCRs and those, like CCK1R that preferentially engage other G protein subfamilies; these diverse modes of binding contribute to the difficulties in using bioinformatic and computational approaches to predict G protein selectivity. Further investigation will be required to understand GPCR-Gs interactions that can trigger destabilisation of the Gs α5 helix “hook” conformation that enables novel modes of engagement with receptors; this will be necessary for prediction of functional Gs binding to GPCRs where limited outward movement of TM6 is an intrinsic structural feature.

**Fig. 5.**
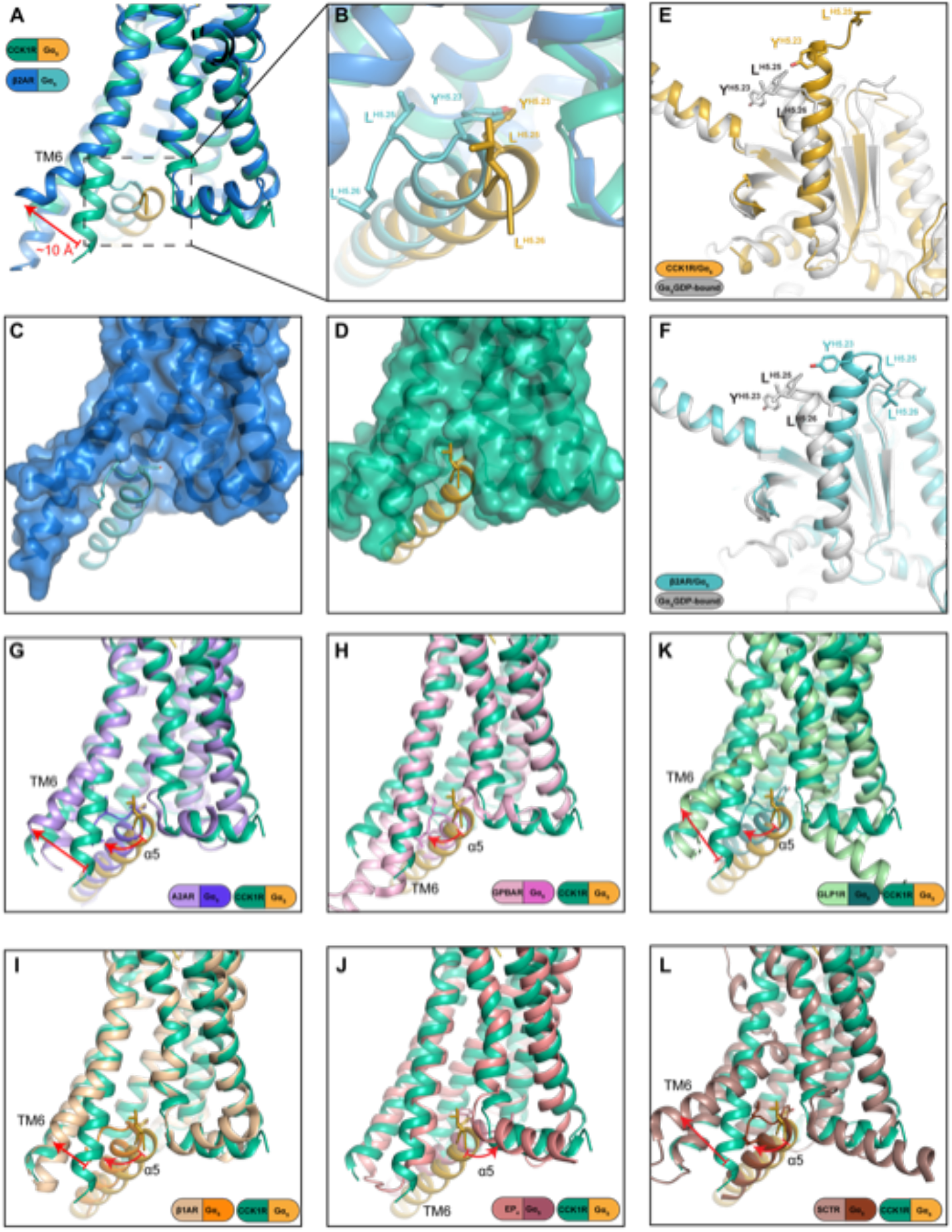
Gαs adopts a C-terminally extended conformation of the α5 helix to enable binding to CCK1R. [**A-D**]. The far C-terminus of the α5 helix of Gαs forms a bulky “hook” conformation for binding to the prototypical Gs-coupled β2-AR [**A-C**] and occupies a large intracellular binding cavity relative to Gs-bound to the CCK1R that has a narrower binding cavity with binding facilitated by formation of an extended conformation of the Gαs C-terminus [**A, B, D**]. In the inactive Gαs-GDP, the Gs C-terminus exhibits the “hook” conformation indicating that this is the principal ground state conformation [**E, F**]. [**G-L**] comparison of the conformation and orientation of Gαs when bound to CCK1R relative to class A, A2AR [**G**, PDB:6GDG], GPBAR [**H**, PDB:7CFM], β1-AR [**I**, PDB:7JJO] and EP4R [**J**, PDB:7D7M], or class B, GLP-1R [**K**, PDB:6X18] and secretin receptor (SCTR) [**L**, PDB:6WZG] that couple predominantly to Gs. Overall, the receptors engage Gs via wide outward movement of TM6 to accommodate the C-terminal hook motif. However, the GPBAR interacts with Gs in a narrower cavity through shallower engagement [**H**], while the EP4R also has a smaller intracellular pocket where Gs engagement is facilitated by unwinding of the far C-terminus of the α5 helix. However, unlike CCK1R, the sidechains are directed into the junction of the base of TM7 and TM1.

Currently, only one other GPCR has been solved with 2 different G protein subfamily transducers; the GCGR bound to Gs or Gi (*14*). For this receptor, the backbone conformation of the receptor was equivalent, regardless of the bound G protein. Here, the default conformation was the prototypical open pocket that facilitates Gs binding with the less bulky Gi1 protein binding into the same pocket with fewer interactions (*14*). In the current study, the CCK1R also exhibited the same backbone conformation, regardless of the bound G protein, but instead this was enabled by conformational change within the Gs protein that allowed it to bind into the smaller CCK1R intracellular binding pocket. While much more work is required, the current data support the contention that the intrinsic conformational flexibility/stability of the receptor (*9, 45*) is a critical feature in modes of G protein coupling for both primary and secondary transducers. The current work also provides supporting evidence for conformational flexibility of the G protein α5 helix as a mechanism for promiscuous coupling.

#### Conclusion

There is much interest in the mechanistic basis for how individual GPCRs bind to different G protein subfamilies and in the features that allow both selectivity and promiscuity in G protein engagement and activation. Herein we demonstrated that the efficiency and selectivity of G protein coupling could be markedly altered through alterations to the availability of different transducers, as well as by plasma membrane cholesterol (**Fig. 1**). In particular, the efficiency of Gs coupling was markedly augmented by either high cholesterol or absence of the primary (Gq/11) transducers demonstrating that the local environment of the receptor within a cell is also critical for the mode of transducer engagement. Differential transducer engagement has been reported for different cellular compartments, linked to different receptor conformations (*46*) and our current observations expand the potential mechanisms for such observations. Our work supports a model for G protein interaction where the extent of GPCR conformational change that occurs upon agonist activation is intrinsic to the individual receptor, with alterations to G protein preference by biased agonists or the local receptor environment more likely to be governed by changes to the conformational dynamics within this primary activated state, rather than large changes to backbone conformations (e.g. larger or smaller intracellular pockets) of the receptor.

## Materials and Methods

### Pharmacological studies

#### Materials

Peptide ligands, natural CCK-26-33 (CCK-8) and an analog for radiolabeling, *D*-Tyr-Gly-[(Nle^28,31^) CCK-26-33], were synthesized in-house as previously described (*47*). LANCE cAMP kit and sodium ^125^Iodine were from Perkin Elmer Health Sciences Inc (Shelton, CT). Fluo-8-AM was from AAT Bioquest (Sunnyvale, CA). Fetal clone II was from Hyclone laboratories (Logan, UT) and Dulbecco’s modified Eagle’s medium, glutamine, zeocin, and soybean trypsin inhibitor were from Thermo Fisher scientific (Waltham, MA). Methyl-β cyclodextrin, 3-isobutyl-1-methyl xanthine and poly-lysine were from Sigma Aldrich (St Louis, MO). All other reagents were of analytical grade.

#### Cell lines

The HEK-293(S) parental cell line and its G protein (Gq, Gs) knock out lines (GsKO, GqKO) generated using the CRISPR-Cas9 approach (*48, 49*) were kindly provided by Dr. Asuka Inoue (Tohoku University, Miyagi, Japan). Each cell line was transfected with human type 1 CCK receptor using PEI (*50*), with clonal cell lines expressing similar levels of receptor selected using CCK-like radioligand binding (*25*). Cells were grown in Dulbecco’s Modified Eagle’s Medium supplemented with 10% Fetal Clone II in a humidified atmosphere containing 5% CO_2_ at 37 °C, and were passaged approximately twice a week.

#### Cholesterol modifications

Cells had their cholesterol composition enhanced using methyl-β-cyclodextrin (MβCD)-cholesterol inclusion complex (*51*).

#### Receptor binding assays

CCK-like radioligand binding assays were performed with intact cells grown in poly-lysine coated 24 -well tissue culture plates, as described (*52*). Non-specific binding was determined in the presence of 1 μM unlabeled CCK. All assays were performed in duplicate and repeated at least three times in independent experiments. Competition-binding curves were analyzed and plotted using the non-linear regression analysis program in the Prism software suite v8.02. (GraphPad, San Diego, CA).

#### Intracellular calcium response assays

CCK-stimulated intracellular calcium responses were quantified in intact cells using Fluo-8-AM using a FlexStation (Molecular Devices, Sunnyvale, CA), as described (*52*). All assays were performed in duplicate and repeated at least three times in independent experiments. Calcium response curves were analyzed and plotted as percentages of the maximal stimulation by 100 μM ATP using non-linear regression analysis in the Prism software suite v8.02.

#### Intracellular cAMP response assays

CCK-stimulated intracellular cAMP responses were measured in intact cells in 384-well white Optiplates using LANCE cAMP kits, as described (*53*). Assays were performed in duplicate and repeated at least three times in independent experiments. The cAMP concentration-response curves were analyzed and plotted using the non-linear regression analysis in Prism v8.02.

#### Statistical comparisons

Comparisons of experimental groups were assessed using analysis of variance (ANOVA) followed by Dunnett’s post-test, or using the Mann-Whitney test, as provided in Prism 8.0 (GraphPad, San Diego, CA). The threshold for statistical significance was set at p < 0.05.

### Structure determination

#### Constructs

Human WT CCK1R was modified to include an N-terminal hemagglutinin (HA) signal sequence, FLAG-tag, 22 amino acids of the N-terminus of the M4 muscarinic receptor to improve expression (*28, 29*), a 3C-protease recognition sequence to remove the N-terminal modifications and a C-terminal 8xhistadine (8xHis) tag (**Fig. S1A**). This construct was cloned in to the pFastBac vector for insect cell expression and pcDNA3.1 for mammalian expression. A construct without modifications was also cloned into pcDNA3.1 for construct validation. Previously described constructs were used to express the heteromeric G protein in insect cells, including a dominant negative form of GαS (DNGαS) and a dual expressing vector containing Gγ2 and 8xHis tagged Gβ1 (*8, 54*). 8xHis tagged Nb35 in pET20b was obtained from B. Kobilka (*39*). For the CCK1R/mGsqi complex, the CCK1R was modified to include the HA signal sequence, N-terminal FLAG-tag, and C-terminal 3C-protease recognition site, 8xHis-tag and mGsqi (**Fig. S1B**; *55*).

#### Insect cell expression

CCK1R, DNGαS, Gβ_1_γ_2_ were co-expressed in the Tni insect cell line (Expression systems) using the Bac-to-bac baculovirus system (Invitrogen). The Tni insect cells were grown to a density of 4 million cells/ml in ESF 921 serum free media (Expression systems) before infection with a 4:2:1 ratio of CCK1R:DNGαS:Gβ_1_γ_2_ baculoviruses. Insect cells were harvested by centrifugation at ~48 hours post infection and cell pellets stored at −80 °C. For the CCK1R/miniGsqi similar methods were, except a 4:1 ratio of CCK1R-mGsqi:Gβ_1_γ_2_ baculoviruses was used.

#### Nb35 and ScFv16 expression and purification

Nb35 was expressed in the periplasm of BL21(DE3) Rosetta E. coli cell line using an autoinduction method (*56*). Transformed cells were grown at 37 °C in a modified ZY media (50 mM Phosphate buffer pH 7.2, 2% tryptone, 0.5 % yeast extract, 0.5% NaCl, 0.6% glycerol, 0.05% glucose and 0.2 % lactose), in the presence of 100 μg/ml carbenicillin and 35 μg/ml chloramphenicol. At an OD_600_ of 0.7 the temperature was changed to 20 °C for ~ 16 hours before harvesting by centrifugation and storage at −80 °C. NB35 was purified as described previously by extracting the periplasm supernatant and Ni-affinity chromatography (*57*).

ScFv16 was expressed in the Tni insect cell line as an excreted product using baculovirus. Media was harvested by centrifugation 72 hours post infection and chelating agents quenched by the addition of 5 mM CaCl2. The media was separated by precipitation by centrifugation and batch bound to EDTA-resistant Ni-Sepharose resin. The column was washed with a high salt buffer (20 mM HEPES pH 7.2, 500 mM NaCl and 20 mM imidazole), followed by a low salt buffer (20 mM HEPES pH 7.2, 500 mM NaCl and 20 mM imidazole), before elution in 20 mM Hepes pH 7.5, 100 mM NaCl and 250 mM imidazole. The eluted protein was dialysed to 20 mM Hepes pH 7.5 and 100 mM NaCl before storage at −80 °C.

#### CCK1R/CCK-8/DNGαS complex purification

Insect cell pellet from 1 L of culture was thawed and solubilised in 20 mM HEPES pH 7.4, 100 mM NaCl, 5 mM MgCl_2_, 5 mM CaCl_2_, 0.5% lauryl maltose neopentyl glycol (LMNG, Anatrace), 0.03% cholesteryl hemisuccinate (CHS, Anatrace) and a cOmplete Protease Inhibitor Cocktail tables (Roche). The resuspended pellet was homogenised in a Dounce homogeniser and complex formation initiated by the addition of 10 μM CCK-8, 5 μg/ml NB35 and 25 mU/ml apyrase (NEB). The solubilisation was incubated stirring at °C for 2 hours before insoluble material was removed by centrifugation at 30,000g for 30 min. The solubilised complex was batch bound to equilibrated M1 anti-Flag affinity resin, rotating at 4 °C for 1 hour. The resin was packed into a glass column and washed with 20 column volumes of 20 mM HEPES pH 7.4, 100 mM NaCl, 5 mM MgCl_2_, 0.01% LMNG, 0.0006% CHS and 1 μM CCK-8 in the presence of 5 mM CaCl_2_, before elution in the same buffer with 5 mM EGTA and 0.1 mg/ml Flag peptide. Eluted complex was concentrated to less than 500 μL in an Amicon Ultra-15 100 kDa molecular weight cut-off centrifugal filter unit (Millipore) and further purified by size exclusion chromatography (SEC) on a Superdex 200 Increase 10/300 GL (Cytiva) with 20 mM HEPES pH 7.4, 100 mM NaCl, 5 mM MgCl_2_, 0.01% LMNG, 0.0006% CHS and 1 μM CCK-8. Eluted fractions were containing complex were pooled and concentrated to 4.1 mg/ml before being flash frozen in liquid nitrogen and stored at −80 °C.

#### CCK1R/CCK-8/mGsqi complex purification

Insect cell pellet from 1.5 L of culture was thawed and resuspended in a hypotonic solution of 20 mM Hepes pH 7.4, 5 mM MgCl_2_ and 1 μM CCK-8. The cells were stirred at room temperature for 15 min to allow lysis to occur. Cell pellet was collected by centrifugation and solubilised in 20 mM HEPES pH 7.4, 100 mM NaCl, 5 mM MgCl_2_, mM CaCl_2_, 0.5% lauryl maltose neopentyl glycol (LMNG, Anatrace), 0.03% cholesteryl hemisuccinate (CHS, Anatrace) and a cOmplete Protease Inhibitor Cocktail tables (Roche). The resuspended pellet was homogenised in a Dounce homogeniser and complex formation initiated by the addition of 10 μM CCK-8, 5 μg/ml ScFv16 and 25 mU/ml apyrase (NEB). The solubilisation was incubated stirring at 4 °C for 2 hours before insoluble material was removed by centrifugation at 30,000g for 30 min. The solubilised complex was batch bound to equilibrated M1 anti-Flag affinity resin, rotating at 4 °C for 1 hour. The resin was packed into a glass column and washed with 20 column volumes of 20 mM HEPES pH 7.4, 100 mM NaCl, 5 mM MgCl_2_, 0.01% LMNG, 0.0006% CHS and 1 μM CCK-8 in the presence of 5 mM CaCl_2_, before elution in the same buffer with 5 mM EGTA and 0.1 mg/ml Flag peptide. Eluted complex was concentrated to less than 500 μL in an Amicon Ultra-15 100 kDa molecular weight cut-off centrifugal filter unit (Millipore) and further purified by size exclusion chromatography (SEC) on a Superdex 200 Increase 10/300 GL (Cytiva) with 20 mM HEPES pH 7.4, 100 mM NaCl, 5 mM MgCl_2_, 0.01% LMNG, 0.0006% CHS and 1 μM CCK-8. The sample was then cleaved with 3C-protease at room temperature for 1 hour and reapplied to SEC. Eluted samples were pooled and concentrated to 4 mg/ml before being flash frozen and stored at −80 °C.

#### SDS-PAGE and western blot

Purified samples were analysed by SDS-PAGE and western blot. Samples were applied to precast TGX gels (BioRad) before staining with InstantBlue Coomassie stain (Sigma-Aldrich) or immediately transferred to PVDF membrane (BioRad) for western blot analysis. Western blots were stained with primary rabbit anti-GNAS (Novus Biologicals, NBP1-31730), primary mouse anti-His antibody (QIAGEN, 34660), secondary goat anti-rabbit 800CW antibody (LI-COR, 926-32211), secondary anti-mouse 680CW antibody (LI-COR, 926-68070) and an in house made Alexa Fluor 488 conjugated mouse anti-flag antibody. Western blots were imaged on a Typhoon 5 imaging system (Amersham).

#### Vitrified sample preparation and data collection

Samples (3 μL) were applied to a glow-discharged UltrAufoil R1.2/1.3 300 mesh holey grid (Quantifoil GmbH, Großlöbichau, Germany) and were flash frozen in liquid ethane using the Vitrobot mark IV (Thermo Fisher Scientific, Waltham, Massachusetts, USA) set at 100% humidity and 4 °C for the prep chamber with a blot time of 10s. Data were collected on Titan Krios microscope (Thermo Fisher Scientific) operated at an accelerating voltage of 300 kV with a 50 μm C2 aperture with no objective aperture inserted and at an indicated magnification of 130kX in nanoprobe TEM mode. A Gatan K3 direct electron detector positioned post a Gatan BioQuantum energy filter (Gatan, Pleasanton, California, USA), operated in a zero-energy-loss mode with a slit width of 15 eV was used to acquire dose fractionated images of the samples. Movies were recorded in hardware-binned mode (previously called counted mode on the K2 camera) with the experimental parameters listed in **Table S1** using 18-position beam-image shift acquisition pattern by custom scripts in SerialEM (*58*).

### Data processing

#### CCK1R/DNGs/CCK-8

7146 micrographs were motion corrected using UCSF MotionCor2 (*59*) and dose weighted averages had their CTF parameters estimated using CTFFIND 4.1.8 (*60*). Particles were picked using the crYOLO software package (*61*) on a pretrained set of weights for GPCRs yielding 6.4 M particle positions. These particles were extracted from the micrographs and subjected to 2D classification and ab initio 3D and 3D refinement in the cryoSPARC (v3.1) software package (*62*) which resulted in a homogeneous well centered particle stack containing 1 M particles. These were then fed into the RELION (v 3.1) software package for further rounds of 2D and 3D classification which led to 800 k particles for initial 3D refinement in RELION. These particles where then polished in RELION and underwent a further round of 3D classification and CTF envelope fitting. A final consensus 3D refinement using 643k particles was performed in RELION the maximization step was carried out using the SIDESPLITTER algorithm (*63*) to yield a final resolution of 1.95 Å (FSC=0.143, gold standard) for the consensus map. Further receptor focused refinements were performed using a mask generated from an initial PDB model and searching a local 1.8 degree Euler angle space.

#### CCK1R/mGsqi/CCK-8

7182 micrographs were motion corrected using UCSF Motioncor2 and dose weighted averages had their CTF parameters estimated using CTFFIND 4.1.8. Particles were picked using the automated template picking routine in RELION 3.1. 2.3 M particles were extracted and cryoSPARC was employed to perform 2D classification, ab initio 3D model generation and initial 3D refinement. The resulting particle stack contained 441 k particles which were then polished in RELION 3.1. The polished particle stack was then fed back into the cryoSPARC software package for a non-uniform 3D consensus refinement and CTF envelope fitting which yielded a 2.44 Å resolution map (FSC=0.143, gold standard). Due to a large amount of conformational flexibility between the receptor and G-proteins, further local refinements in cryoSPARC were used to calculate high quality maps of either the CCK1 receptor (2.57 Å) or the mGsqi G-protein complex (2.43 Å) which were used to generate a PDB model.

#### CCK1R/CCK-8/DNGs model building

An initial homology model of CCK1R was created using SWISS-MODEL (*64*) using the active structure of the serotonin 5-HT1B receptor as a template (PDB ID: 6G79). The CCK1R model was placed into the receptor focused cryo-EM map by the MDFF routine in namd2 (*65*). The CCK-8 peptide was manually built using COOT (*66*), the sulfotyrosine residue was imported from the monomer library (TYS) and geometry restrains generated with eLBOW within PHENIX (*67*). The model was refined with repeated rounds of manual model building in COOT and real-space refinement within PHENIX (*68*). The G protein (DNGαS/Gβ1/Gγ2) and NB35 from the structure of GLP-1 receptor-Gs complex (PDB ID: 6B3J) was rigid body placed into the consensus cryo-EM map using the map fitting tool of UCSF ChimeraX (*69*). The G protein was further refined to the consensus cryo-EM map with repeated rounds of manual model building in COOT and real-space refinement within PHENIX. Lastly the CCK1R/CCK-8 model was combined with the G protein and real-space refined to the consensus cryo-EM map using PHENIX. The model quality was assessed using MolProbity (*70*) before PDB deposition.

#### CCK1R/CCK-8/mGsqi model building

The higher resolution model of CCK1R/CCK-8 obtained from the GαS structure was rigid-body placed a receptor focused map cryo-EM map using ChimeraX. The model was refined with repeated rounds of manual model building in COOT and real-space refinement within PHENIX. The β1 and γ2 were obtained from the higher resolution GαS structure, the miniGsqi was modified from the OX_2_R structure (PDB ID: 7L1U) and the subunits were rigid-body placed in the G protein focused map. The model was refined with repeated rounds of manual model building in COOT and real-space refinement within PHENIX (Afonine et al, 2018). Lastly the CCK1R/CCK-8 model was combined with the G protein and real-space refined to the consensus cryo-EM map using PHENIX. The model quality was assessed using MolProbity before PDB deposition. The scFv16 was not modelled due to poor side chain density.

#### Model interaction analysis

Interactions and hydrogen bonds were analysed using UCSF chimeraX package, with relaxed distance and angle criteria (0.4 Å and 20° tolerance, respectively). The interfaces were further analysed using PDBePISA (*71*). Figures were generated using UCSF chimeraX and PyMOL Molecular Graphics System, Version 2.3 (Schrödinger, LLC).

#### 3D variability analysis (3DVA)

3DVA was performed using the cryoSPARC software package, based on the consensus refinements of the complexes. For the CCK1R-mGsqi data, six principal components (Comp0 to Comp5) were used, and for the CCK1R-Gs data five principal components (Comp0 to Comp4) were used to analyse the principal motions in the cryoEM data, each separated into 20 frames (frame 0 to 19). Cryosparc 3DVA outputs were used for dynamic analyses of the CCK complexes and visualised using the command *vseries* as implemented in ChimeraX (*69*). The backbones of the consensus refinement models were flexibly fitted into the frame 0 and 19 density maps of the 3DVA principal components using Isolde (*72*), implemented in ChimeraX (*69*). Morphs of models (aligned using the *matchmaker* command, or aligned by densities) were created in Chimera and ChimeraX (*69, 73*). Movie editing was performed using Adobe Premiere Pro 2020.

## Acknowledgments

Molecular graphics and analyses were performed with UCSF ChimeraX, developed by the Resource for Biocomputing, Visualization, and Informatics at the University of California, San Francisco, with support from National Institutes of Health R01-GM129325 and the Office of Cyber Infrastructure and Computational Biology, National Institute of Allergy and Infectious Diseases. The work was supported by the Monash University Ramaciotti Centre for Cryo-Electron Microscopy and the Monash MASSIVE high-performance computing facility.

## Author contributions

J.M. expressed and purified receptor complexes. H.V. performed EM data collection during optimisation of samples. R.D. performed cryo-sample preparation and imaging to acquire high-resolution EM data. M.J.B. processed the EM data and performed EM map calculations, including 3D multivariate analysis. J.M. and M.J.B. built the protein models for consensus maps and performed refinement. K.G.H and X.X. performed pharmacological characterisation experiments. S.G.B.F. supervised experiments for characterisation of expression constructs. S.J.P. and D.M.T. cross-checked the atomic models, and S.J.P. built protein models into the 3D variance data. J.M., M.J.B., S.J.P., K.G.H, L.J.M, D.W. and D.M.T assisted with data interpretation, figure and manuscript preparation. All authors reviewed and edited the manuscript; P.M.S., L.J.M., D.M.T., A.C., D.W. designed the project and/or interpreted data. P.M.S., D.W., D.M.T. and L.J.M. supervised the project. P.M.S., J.M. D.M.T., M.J.B. and L.J.M. generated figures and wrote the manuscript.

## Data and materials availability

All relevant data are available from the authors and/or included in the manuscript or Supplementary Information. Atomic coordinates and the cryo-EM density map have been deposited in the Protein Data Bank (PDB) under accession numbers 7MBX (CCK8:CCK1R:DNGs complex) and 7MBY (CCK8:CCK1R:mGq/s/i complex), and Electron Microscopy Data Bank (EMDB) accession numbers EMD-23749 (Gs complex) and EMD-23750 (Gq complex). All constructs used in the study are available from the authors upon reasonable request.

## Supplementary Materials

**Fig. S1.**
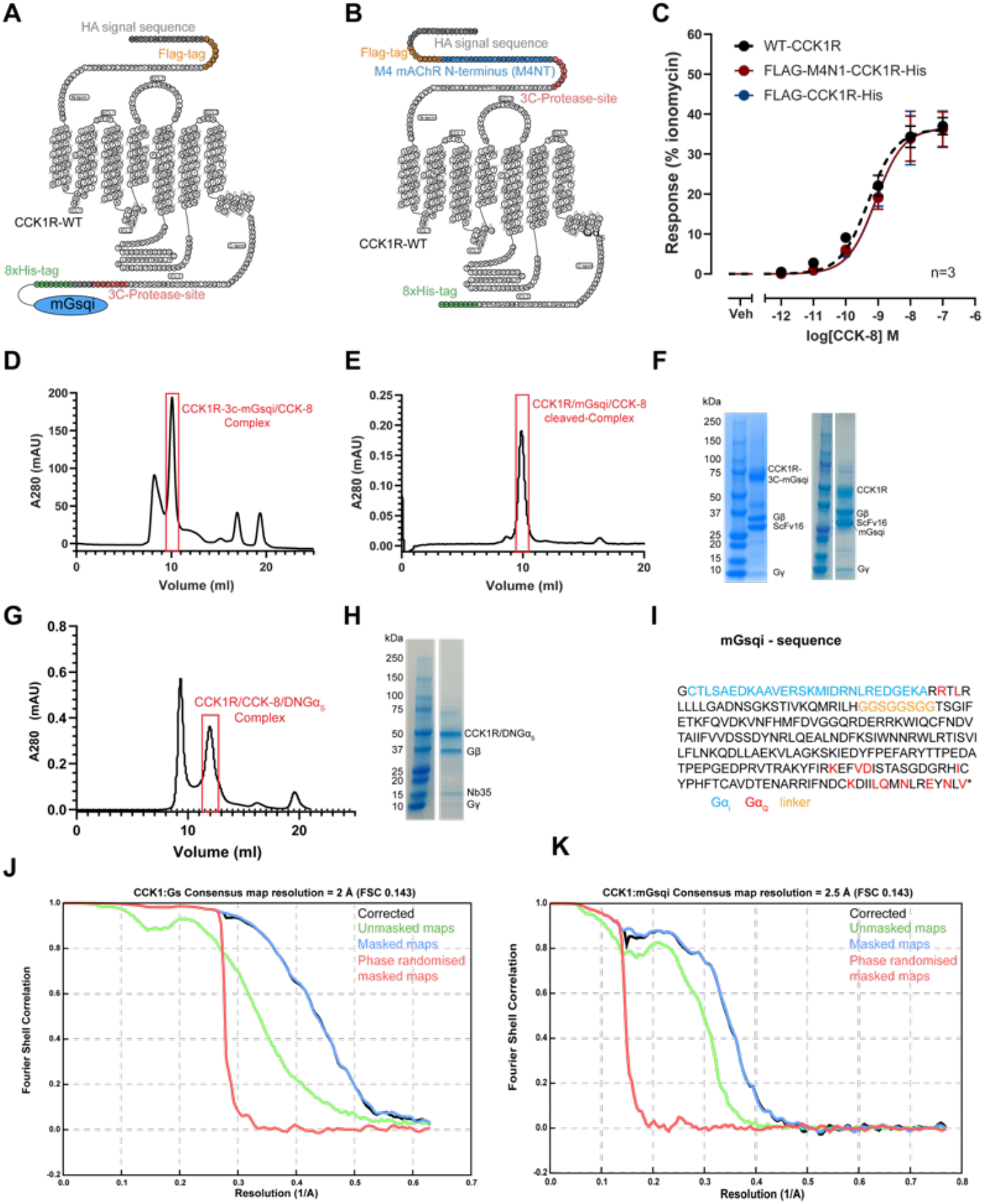
Expression and purification of CCK-8/CCK1R/G protein complexes. [**A, B**] Snake plot of the CCK1R expression constructs for formation of mGsqi [**A**] or Gs [**B**] complexes. The construct for mGsqi complex formation contained an N-terminal Hemagglutinin (HA) signal sequence (grey shading), followed by a Flag-epitope tag (yellow shading), with a 3C protease cleavage site (red shading) inserted at the C-terminus followed by an 8-His tag (green shading) and the mGsqi (blue schematic). For Gs complex formation, the construct contained an N-terminal Hemagglutinin (HA) signal sequence, followed by a Flag-epitope tag, a M4 mAChR N-terminal sequence (blue shading) and a 3C-cleavage site (red sequence) with an 8-His tag fused to the C-terminus. [**C**] The expression constructs (red circles or blue circles) were cloned into a mammalian expression vector and concentration-responses to CCK-8 in an iCa^2+^ mobilisation assay were established relative to WT (black circles). [**D**], **E**] SEC of affinity purified of the CCK-8/CCK1R/mGαsqi/Gβγ1γ2/scFv16 complex before [**D**] and after [**E**] sample cleavage with 3C protease; the peak used for SDS-PAGE analysis is boxed in red. [**F**] Coomassie blue stained SDS-PAGE of the peak sample from [**D**] (left panel) or [**E**] (right panel). [**G**] SEC of affinity purified of the CCK-8/CCK1R/DNGαs/Gβ1γ2/Nb35 complex. [**H**] Coomassie blue stained SDS-PAGE of the peak sample from [**G**]. [**I**] amino acid sequence of the mGsqi chimera illustrating the origin of the different segments. [**J, K**] Gold standard Fourier shell correlation curves for the final map and map validation from half maps showing the overall nominal resolutions of 2.5 Å for the CCK1R-“Gq-mimetic” complex [**J**] and 2.0 Å for the CCK1R-Gs complex [**K**].

**Fig. S2.**
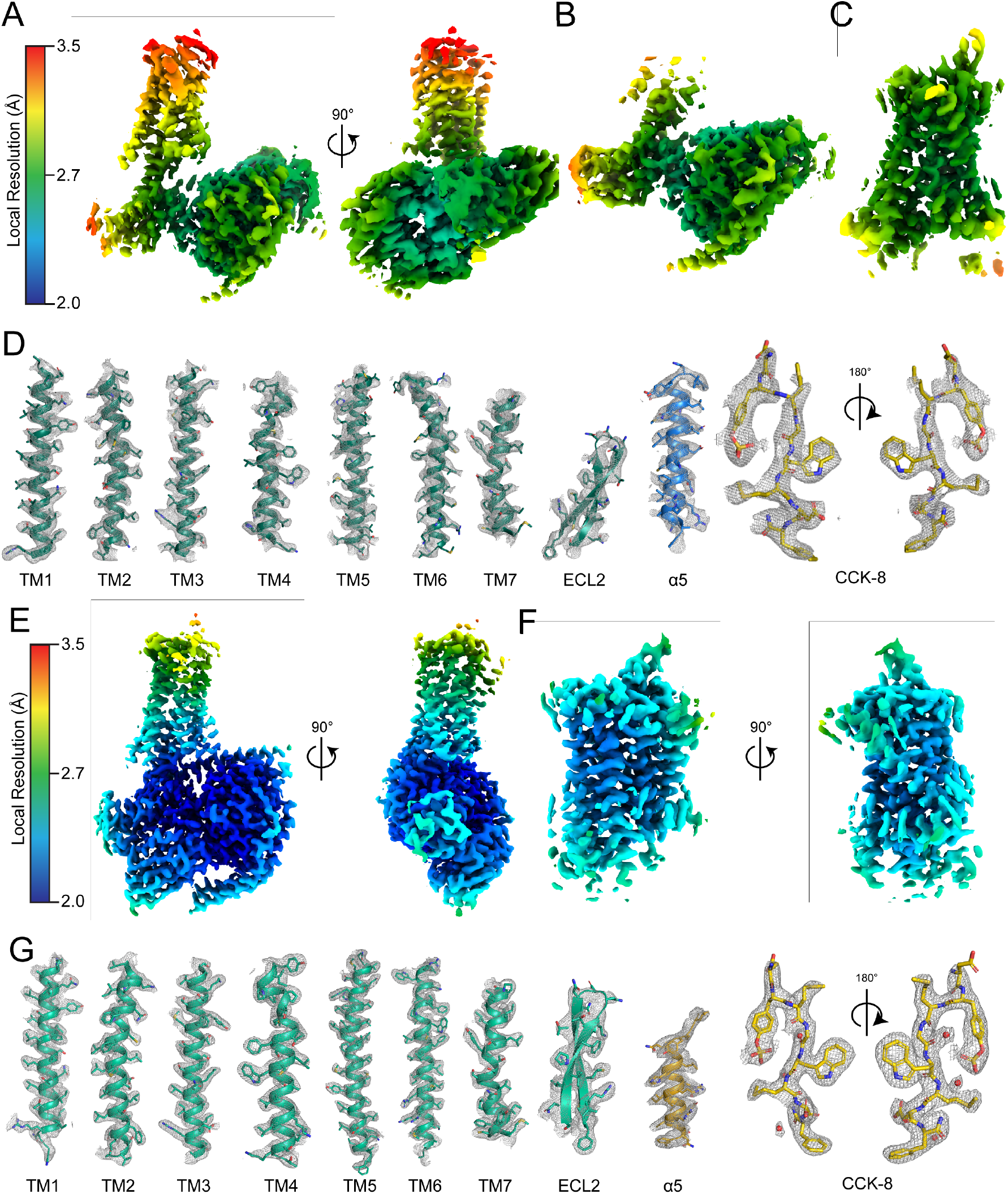
Local resolution and atomic modelling into the cryo-EM density maps. Local resolution of [**A**] the consensus map, [**B**] G protein-focused refinement and [**C**] receptor-focused refinement of the CCK-8/CCK1R/mGαsqi/Gβ1γ2/scFv16 complex. [**D**] Density maps and models are illustrated for all seven transmembrane helices and ECL2 of CCK1R, the α5 helix of the Gα subunit and the CCK-8 peptide. [**E**], [**F**] Local resolution of the consensus [**E**] and receptor-focused [**F**] maps for the CCK-8/CCK1R/DNGαs/Gβ1γ2/Nb35 complex. [**G**] Density maps and models are illustrated for all seven transmembrane helices and ECL2 of CCK1R, the α5 helix of the Gα subunit and the CCK-8 peptide. Protein backbone is displayed in ribbon format with amino acid sidechains in stick representation, coloured by heteroatom. The cryo-EM density was zoned at 1.8 Å.

**Fig. S3.**
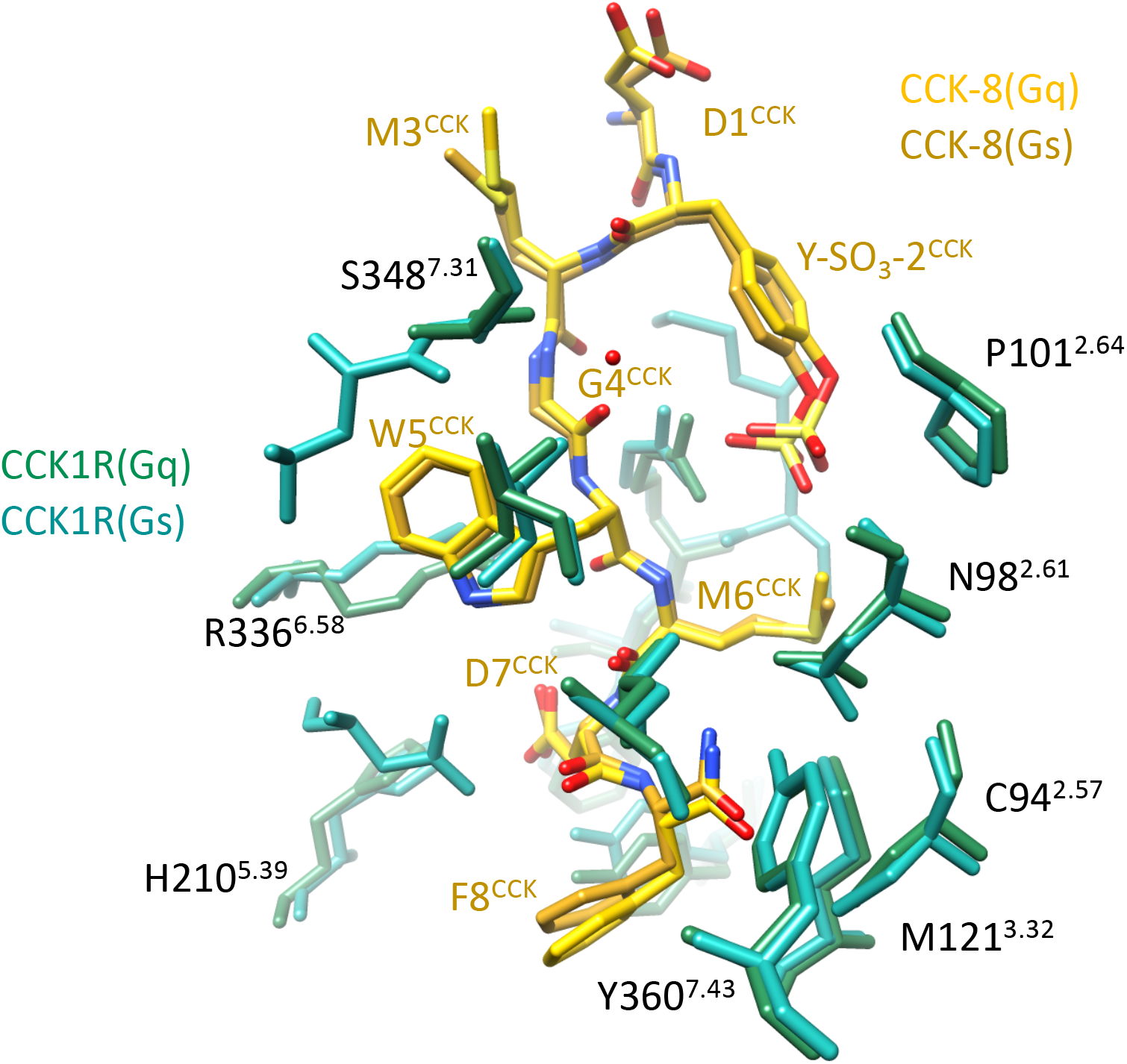
CCK-8 has an equivalent pose and interaction with CCK1R in both the Gs and Gq-mimetic protein complexes. The CCK-8 residues are displayed in stick format, coloured by heteroatom (Gs-complex, gold; Gq-mimetic complex, yellow). CCK1R sidechains that interact with the peptide are shown in stick format (Gs-complex, light green; Gq-mimetic complex, dark green).

**Fig. S4.**
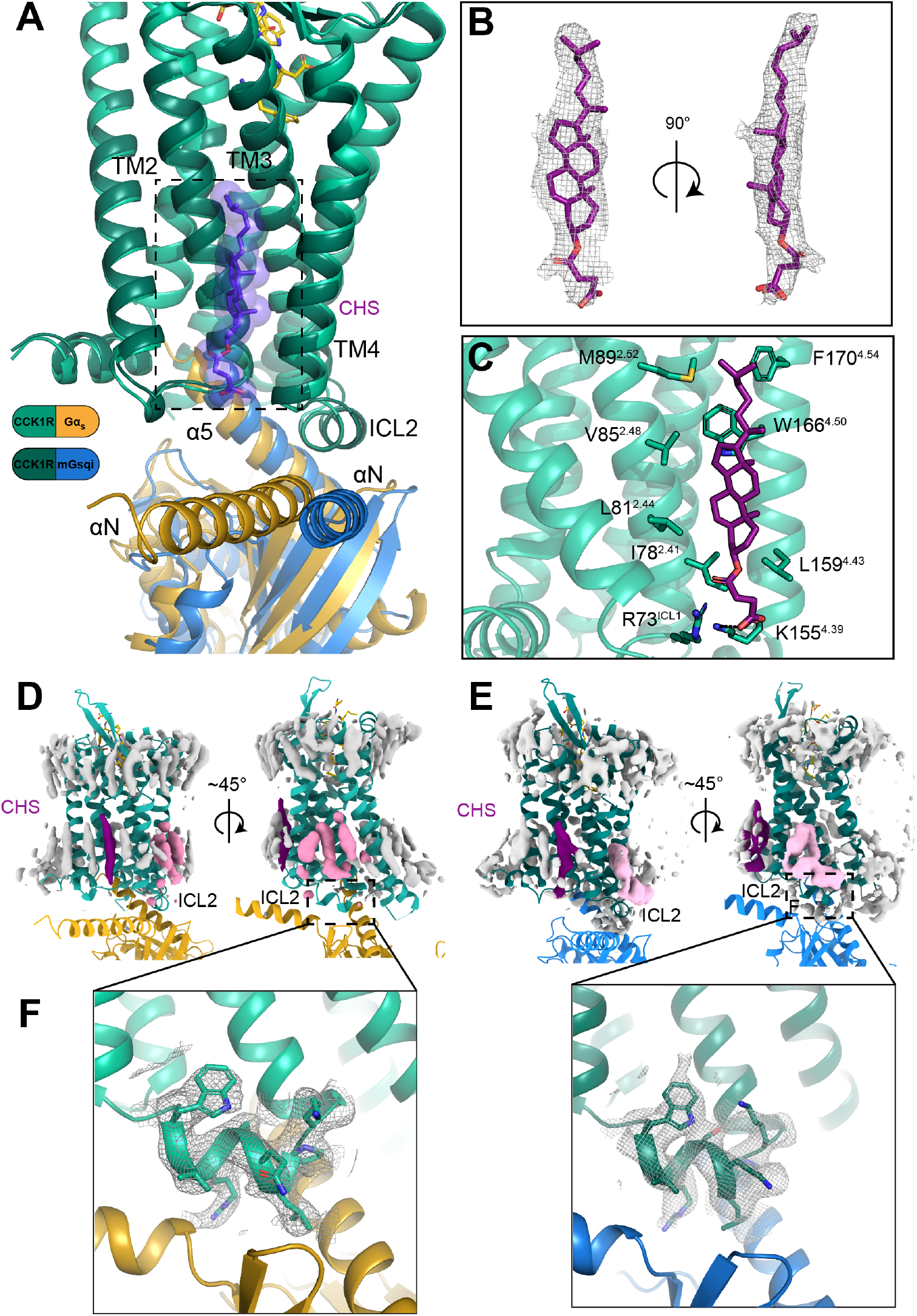
The active CCK1R is surrounded by annular lipids. [**A**] Alignment of the Gs and mGsqi structures illustrating the location of the Gαs protein and modelled lipid. [**B**] Model of the cholesteryl hemi succinate (CHS) in the cryo-EM density. [**C**] The modelled CHS interacts with TM2 and TM4 of CCK1R. [**D-E**] Receptor-focused maps for the Gs protein-complex [**D**] and Gq-mimetic protein complex [**E**] are shown, coloured by component (CCK1R, green; CCK-8, yellow; Gαs protein, gold; Gαq-mimetic protein, blue; unmodelled lipids, grey; putative CHS, purple). There is weak cryo-EM density in the predicted binding site for the allosteric cholesterol abutting ICL2 (pink density). [**F**] The cryo-EM density of ICL2 (zoned at 1.8 Å) in the Gs-complex supports a single predominant conformation, while the density in the Gq-mimetic complex is less well resolved.

**Fig. S5.**
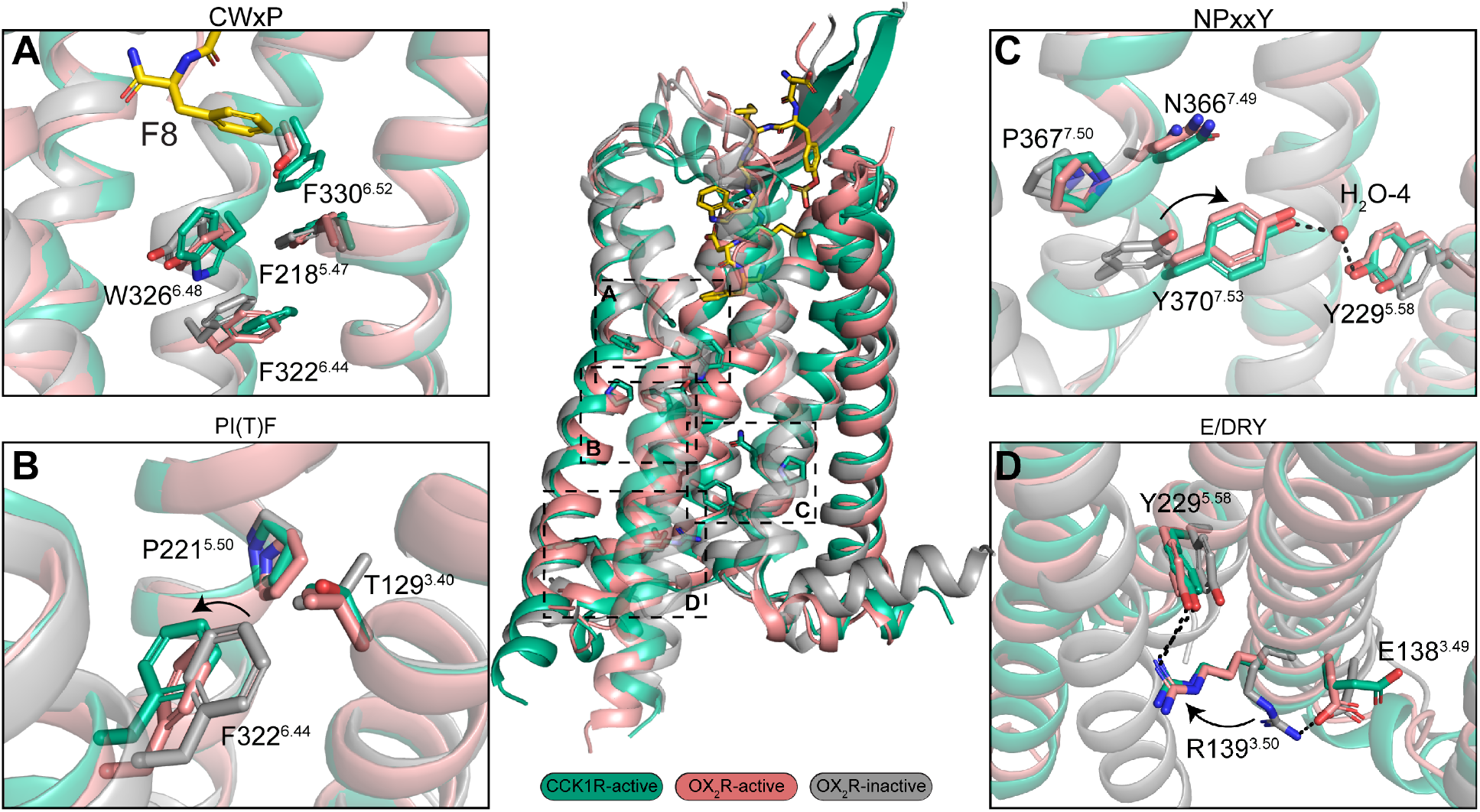
Comparison of activation motifs between CCK1-active (green) and OX_2_R-active (pink) and inactive (grey). [**A**] CWxP motif. While there is no direct interaction between CCK-8 and W326^6.48^ of the CWxP motif, residue F8^CCK^ interacts with F330^6.52^, which in turn interacts with W326^6.48^. Of note, residue 6.48 is not conserved in OX_2_R and there is no movement between the inactive and active states at this position. Given the absence an inactive CCK1R structure the role of the CWxP motif in CCK1R is unclear. Nevertheless, the position of W326^6.48^ stabilises the outward rotation of F322^6.44^ within the PI(T)F motif. [**B**] Comparison of the PI(T)F motif shows this activation motif is largely conserved between CCK1R and OX_2_R. Upon activation there is a modest movement of F^6.44^ for OX_2_R. CCK1R is even further shifted away from T129^3.40^ and P221^5.50^. [**C**] NPxxY motif. Upon activation there is a clear rotation of Y^7.53^ for OX_2_R. CCK1R Y370^7.53^ overlays with the active form of OX_2_R and forms a water mediated interaction Y229^5.58^. [**D**] E/DRY motif. In the OX_2_R inactive state R^3.50^ forms a salt-bridge with E^3.49^; these residues are conserved in CCK1R. Upon activation R^3.50^ rotates up and inwards to form a hydrogen bond with Y^5.58^, the position of these residues overlay with the active CCK1R.

**Fig. S6.**
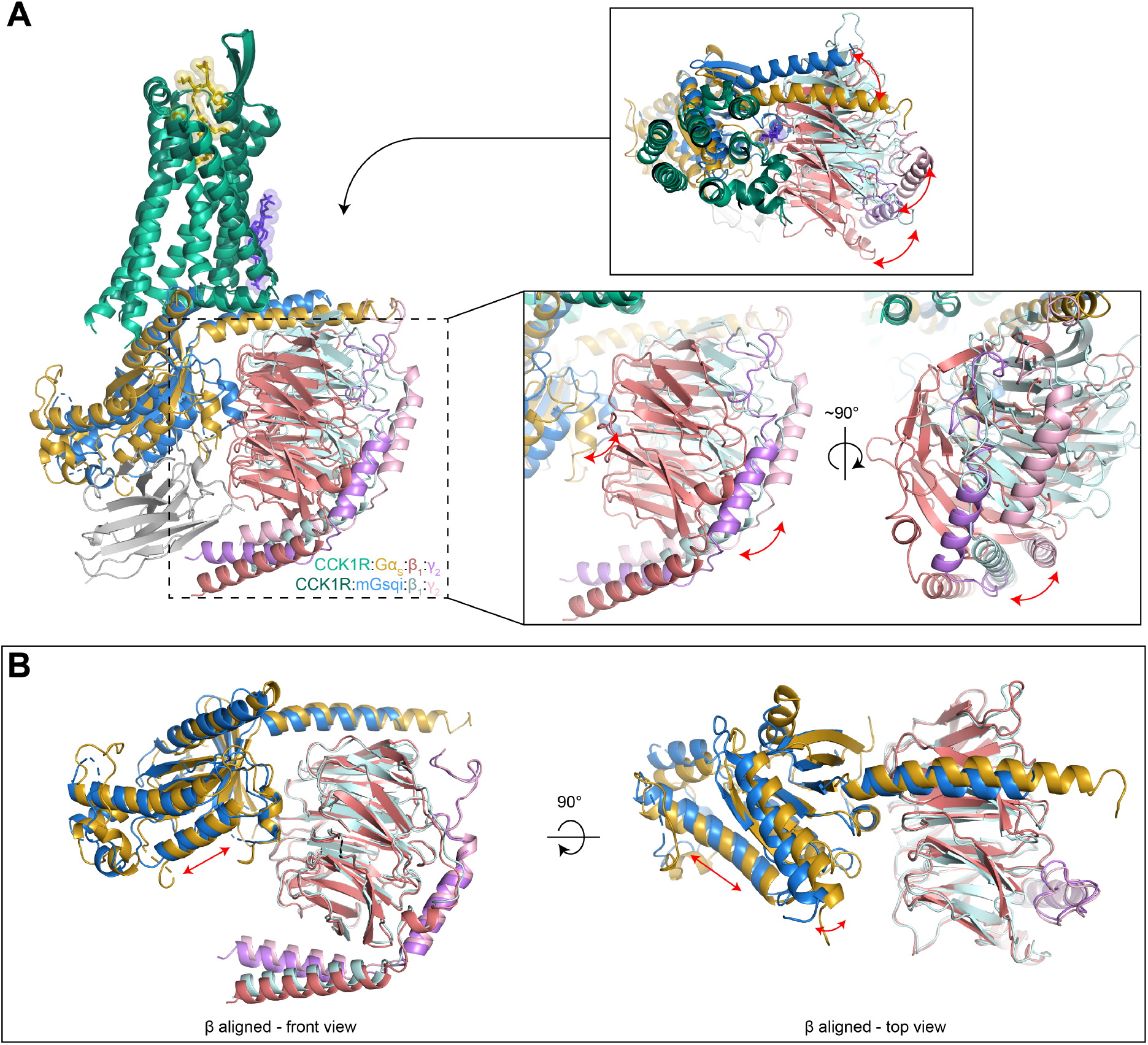
The Gs and Gq-mimetic proteins have distinct orientations in the consensus structures. [**A**] Orientation of the G proteins following alignment on CCK1R. The protein backbone is displayed in ribbon format coloured according to the displayed legend. The CCK-8 peptide (yellow; stick and surface representation) and modelled CHS (purple, stick and surface representation) are also shown (main panel). The proteins were aligned on the receptor. Gs and mGsqi have a different angle of engagement and this is propagated to larger changes in the relative positions of the Gβ and Gγ subunits (inset panels). [**B**] Alignment of the Gβ subunits reveals that the Gα subunits have distinct rotational and translational positions in the heterotrimer.

**Fig. S7.**
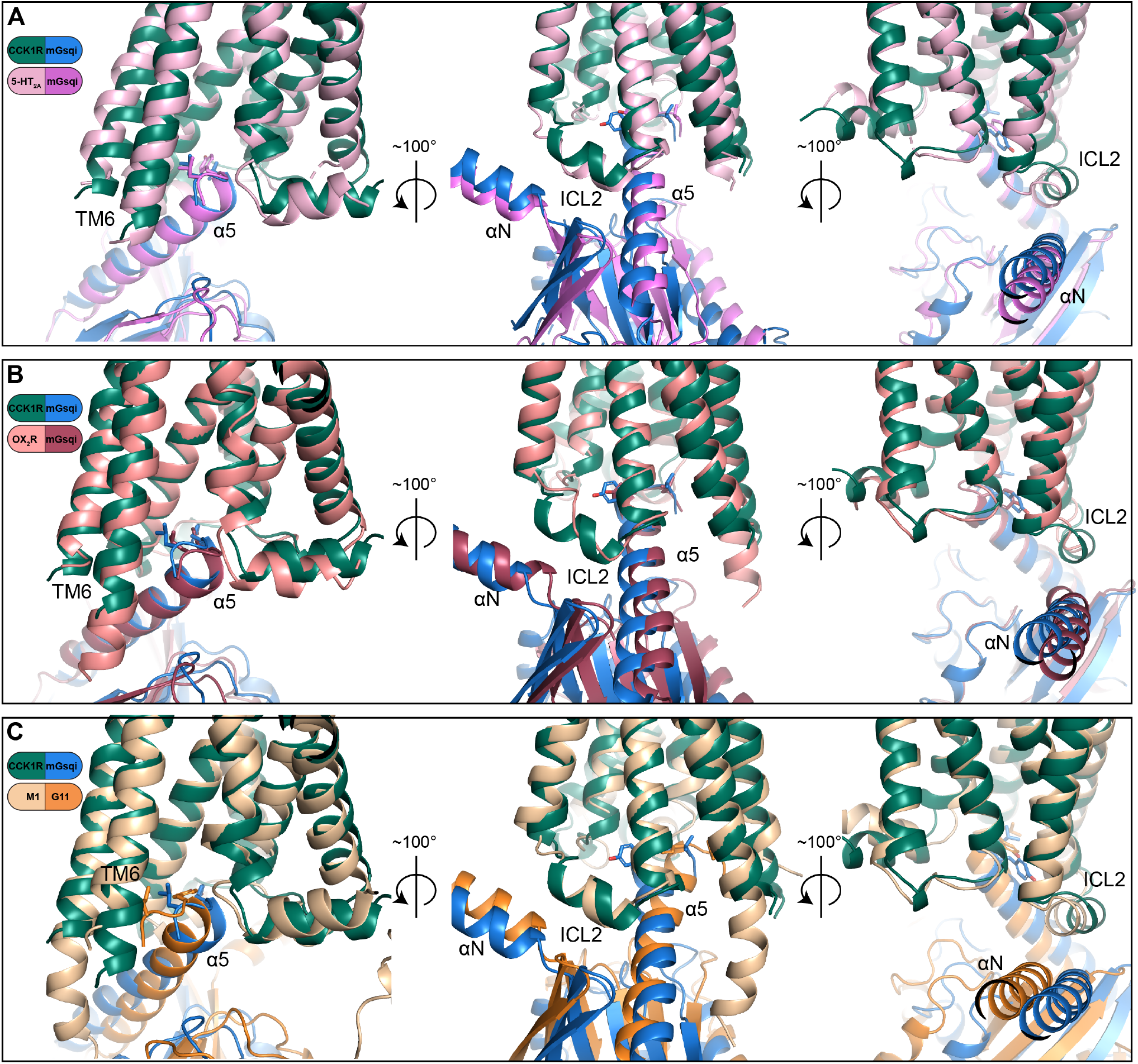
Comparison of the orientation and conformations of Gq-mimetic proteins in class A GPCRs. Each of the available Gq (Gq-mimetic chimera) bound structures is displayed relative to the structure of the CCK1R-mGsqi complex. [**A**] CCK1R-mGsqi and 5HT_2A_R-mGsqi (PDB:6WHA). [**B**] CCK1R-mGsqi and OX_2_R-mGsqi (PDB:7L1U). [C] CCK1R-mGsqi and M1 mAChR-G11/i chimera [PDB:6OIJ). The structures are aligned to the CCK1R. The protein backbone is shown in ribbon format coloured according to the displayed legends. The C-terminal residues of the G protein α5 helix are shown in stick representation coloured by heteroatom. All receptors have a similar, narrow intracellular G protein binding cavity with only small differences in the angle or translational position of the α5 helix when bound to the receptor [**A-C**], with greatest difference seen between CCK1R and M_1_ mAChR complexes [**C**].

**Table S1:** Data collection and refinement statistics.

| **Data Collection** | **CCK1R/Gs/CCK-8** | **CCK1R/mGsqi/CCK-8** |
| --- | --- | --- |
| Micrographs | 7146 | 7182 |
| Electron dose (e<sup>-</sup>/Å<sup>2</sup>) | 70.56 | 63.9 |
| Voltage (kV) | 300 | 300 |
| Pixel size (Å) | 0.65 | 0.65 |
| Defocus range (μm) | 0.5-1.5 | 0.5-1.5 |
| Symmetry imposed | C1 | C1 |
| Particles (final map) | 643k | 444k |
| Resolution (0.143 FSC) (Å) | 1.95 | 2.44 |
| **Refinement** |  |  |
| CC<sub>map_model</sub> | 0.72 | 0.68 |
| Map sharpening B factor (Å<sup>2</sup>) | -38.0 | -73.2 |
| **Model Quality** |  |  |
| R.m.s. deviations |  |  |
| Bond length (Å) | 0.004 | 0.005 |
| Bond angles (°) | 0.782 | 0.914 |
| Ramachandran |  |  |
| Favoured (%) | 98.47 | 98.27 |
| Outliers (%) | 0.00 | 0.00 |
| Rotamer outliers (%) | 0.45 | 0.0 |
| C-Beta deviations (%) | 0 | 0 |
| Clashscore | 2.22 | 4.48 |
| MolProbity score | 1.00 | 1.25 |

## Supplementary Video Legends

**Video S1. Morph between the consensus structures of the CCK-8:CCK1R in complex with mGsqi or Gs.** Structures are displayed in ribbon format with the mGsqi-bound complex in blue, Gs-bound complex in yellow. The morphed coordinates are coloured white. The initial transition shows the full complex, the second transition a close up of the receptor-G protein interface focused on the α5 helix of the Gα subunit and the final transition a close up of the receptor-G protein interface focused on the αN-αH5 junction and receptor ICL2. Morphs between conformations were created in Chimera. The stabilising Nb35 and scFv16 have been omitted for clarity.

**Video S2. 3D variance analysis (3DVA) of the cryo-EM data for CCK1R complexes.** In the first transition, the complex with mGsqi was parsed into 6 principal components with motions for each component displayed side-by-side. In the second transition, the complex with Gs was parsed into 5 principal components with individual components displayed side-by-side. For both complexes the receptor and peptide were highly dynamic in one or more of the component trajectories, with the receptor-G protein interface exhibiting greater relative dynamics for the mGsqi complex over the Gs complex. The 3DVA trajectories are displayed suing the ChimeraX *Volume Series* command.

**Video S3. Morph of the start and end frames of component trajectories from 3D variance analysis (3DVA) of the CCK1R-G protein interface.** Maps and models are shown first for the complex with mGsqi followed by the complex with Gs. The first transition illustrates the full complex and region of focus for the subsequent transition that illustrates individual principal components of motion. The modelled protein is displayed in ribbon format and the map in grey transparent surface representation. CCK1R (dark green, mGsqi complex; light green, Gs complex), mGsqi (blue), Gs (gold), Gβ1(light red), Gγ2 (pink) and Nb35 (Gs-complex; grey) are shown.

**Video S4. Progressive morph between the start and end frames of the individual principal components of motion of CCK1R complexes modelled from 3DVA.** Models of the complexes with mGsqi (left panels) or Gs (right panels) are displayed in ribbon format illustrating the CCK1R-G protein interface from either the “front” (focus on Gαhelix 5; upper panels) or “rear” (focus on GαN/ICL2; lower panels). The transitions commence with Frame 0 of the first component (0) then Frame 19 (component 0) proceeding to Frame 0 of the second component (1) then progressively through to the end (Frame 19, component 5 (mGsqi); Frame 19, component 4 (Gs)). The mGsqi cryo-EM data was parsed into 6 components while the Gs cryo-EM data was parsed into 5 components. The data illustrate the extent of conformational dynamics for the two complexes with greater relative motion overall of the CCK1R-mGsqi complex relative to the CCK1R-Gs complex.

**Video S5. Conformational transitions for “activation” of Gαs proteins.** The video displays the different transitions (morph between structures) that Gαs undergoes when moving from the inactive GDP-bound state (PDB: 6EG8) to the G_0_ state (guanine nucleotide free) induced by binding to selected activated class A or class B GPCRs. **Transition 1.** Inactive to CCK1R-bound. **Transition 2.** Inactive to β2-AR-bound. **Transition 3.** β2-AR-bound to CCK1R-bound. **Transition 4.** Inactive to GLP-1R-bound to CCK1R-bound. **Transition 5.** Inactive to EP4R-bound to CCK1R-bound. The protein backbone is displayed in ribbon format with the C-terminal residues of the α5 helix shown also in stick format (coloured by heteroatom). Gs-GDP (grey; PDB: 6EG8), CCK1R-bound (gold), β2-AR-bound (blue; PDB: 3SN6), GLP-1R-bound (green; PDB:6X18), EP4R-bound (red; PDB: 7D7M). The α5 helix has a ‘hook’ conformation in the inactive G protein and this conformation is maintained in most active state structures. In contrast, the far C-terminal residues of the α5 helix unwinds to enable binding to the CCK1R. Unwinding of the α-5 helix is also seen with the EP4R complex, but this is accommodated by projection of the C-terminal amino acid sidechains between the base of TM7 and TM1 (see **Fig. 5J**).

## Supplementary PDB file captions

**PDB S1.** CCK1_receptor_focused_1_real_space_refined-51-dmt-coot-7_NH2.pdb PDB model from the receptor focused refinement of the CCK-8/CCK1R/Gs complex.

**PDB S2.** mGsQi_real_space_refined_003-coot-13_FINALd.pdb PDB model from the G protein focused refinement of the CCK-8/CCK1R/mGsqi complex.

**PDB S3.** CCK1_mGsQ-coot-060323-fromRSR10-coot-13-coot-4a_real_space_refined_009_NH2C.pdb PDB model from the receptor focused refinement of the CCK-8/CCK1R/mGsqi complex.

